# Podoplanin-Linked Mesenchymal Shift Synergizes with CCR7 driving Lymphatic Metastasis and Tumor Progression in Breast Cancer

**DOI:** 10.1101/2025.03.17.643704

**Authors:** Zhi Wang, Lise Martine Ingebriktsen, Tove Bekkhus, Anastasia Bazioti, Wenyang Shi, Joakim Harry Lehrstrand, Michail Angelos Panagias, Martina Vigorelli, Lyusong Ma, Sarah Ring, Elisabeth Wik, Erling Andre Hoivik, Jonas Fuxe, Maria H. Ulvmar

**Author notes:** Correspondence to: MH Ulvmar, Department of Medical Biochemistry and Microbiology, Uppsala University, Uppsala, Sweden. Shared first authors (Zhi Wang, Lise Martine Ingebriktsen, Tove Bekkhus).

## Abstract

Epithelial-to-mesenchymal-transition (EMT) and upregulation of chemokine receptors are linked to lymphatic metastasis in various types of cancer including breast cancer. However, the underlying molecular mechanisms are not fully understood. Using a triple-negative mammary carcinoma model, conditioned for lymphatic metastasis through expression of the C-C chemokine receptor 7 (CCR7), we identified intrinsic tumor plasticity associated with EMT as a critical determinant for effective metastasis. Specifically, tumor cells upregulate Podoplanin (PDPN) as part of a spontaneous mesenchymal shift that promotes lymphatic dissemination and tumor growth. Intriguingly, the expression of CCR7, or PDPN alone, was not sufficient for these effects. CCR7- and PDPN-positive tumors displayed an immune-cold tumor profile, compared to control, characterized by reduced tumor-infiltrating T-cells. Consistent with this, we found that spontaneous upregulation of PDPN was linked to hypoxia and was associated with the downregulation of homeostatic interferon signaling, and increased expression of collagens. Analysis of single-cell data sets showed that PDPN expression was heterogeneous and correlated significantly with EMT markers, collagen expression, and hypoxia hallmark in human breast cancer cell lines and primary human triple negative (TN) breast cancer, mirroring findings in the mouse model. Further analysis of the human breast cancer METABRIC-microarray datasets supported an association between a high CCR7/PDPN mRNA expression score and aggressive breast cancer subtypes with an independent prognostic value in lymph node-positive tumors. Together, these findings highlight a critical role of tumor microenvironment-driven tumor plasticity and molecular synergy between CCR7 and PDPN-coupled EMT shift in promoting chemokine receptor-mediated tumor dissemination and progression.

**Summary:** CCR7-driven lymphatic metastasis requires a mesenchymal shift with PDPN upregulation. This tumor plasticity, linked to hypoxia and immune evasion, suggests CCR7-PDPN synergy in aggressive breast cancer progression and poor prognosis.

## Background

Lymph node (LN) metastasis serves as an independent prognostic indicator in several cancer types, including breast cancer, where it is strongly associated with an increased risk of disease progression and distant metastases (1, 2). Both intrinsic and extrinsic factors contribute to lymphatic metastasis, including epithelial-to-mesenchymal transition (EMT), immune evasion, metabolic reprogramming, and microenvironmental conditioning (3–6). The factors that determine tumor spread and their interactions may vary across different tumor types and stages (7). These factors can also influence subsequent processes, such as the risk of further metastatic spread and activation of tumor immunity.

Tumor cell expression of C-C chemokine receptor 7 (CCR7) has been identified as a tumor cell intrinsic factor that promote lymphatic metastasis, as demonstrated both in human cancers and experimental work (8–12). CCR7 enables tumor cells to exploit chemotactic signaling pathways normally used by immune cells to migrate into lymphatic vessels (13, 14) and subsequently transit into tumor-draining lymph nodes (TDLNs). Upregulation of CCR7 in mammary tumor cells, and its ligand CCL21 in lymphatic endothelial cells, has been linked to TGF-β1-induced EMT and lymphatic metastasis (15). Data from human breast cancer suggest that the effect of tumor-expressed CCR7 on driving lymphatic metastatic spread may be limited by tumor stage or subtype and microenvironmental factors (16, 17), but details are still lacking.

Here we provide new insights into the interplay between CCR7 in lymphatic metastasis promotion and tumor cell plasticity using EO771 (also known as E0771), a model of triple negative (TN) like mammary carcinoma (18, 19). EO771, similar to most carcinoma models of cancer in the C57Bl6 background, display limited lymphatic metastasis (18–20). Our data suggests that a specific shift in the EMT profile, which is linked to increased expression of the glycoprotein podoplanin (PDPN) and modulation of collagen expression in tumor cells, is required for efficient CCR7-driven metastasis.

Importantly, EMT is not a binary switch, but rather a process that can display reversibility and adaption to the microenvironment (21, 22). Epithelial-to-mesenchymal plasticity (EMP) in tumor cells has been linked to adaptation to different growth conditions, metastasis, and immune evasion (21, 22). Our data supports this notion by coupling spontaneous PDPN induction with the microenvironment *in vivo*. Cell-specific changes also include suppression of the interferon response, suggesting an immune evasion phenotype in cells with single PDPN or CCR7 expression, an effect that is further enhanced in cells co-expressing both markers. We also demonstrate that heterogeneity in PDPN expression can occur in human BC, specifically TNBC, where a higher PDPN cell proportion can be linked to a higher EMT score together with gene hallmarks of hypoxia. This indicates that the central molecular effects associated with the upregulation of PDPN in the EO771 mouse model can be translated to human breast cancer. We further link our experimental data to human breast cancer using the METABRIC data gene-expression data set. Taken together, our findings support an impact of the tumor microenvironment and tumor cell plasticity in facilitating chemokine-driven tumor dissemination and progression.

## Results

### Expression of CCR7 in the triple-negative breast cancer cell line EO771 leads to increased tumor growth and lymphatic metastasis *in vivo*

CCR7 expression is known to increase lymphatic metastasis across multiple tumor models and has been associated with lymphatic metastasis in several types of human cancers (23). To condition the mammary carcinoma cell line EO771 for enhanced metastatic potential, expression of the chemokine receptor CCR7 was introduced (Supplementary Fig. 1A) together with tdTomato for the sensitive tracing of tumor cells. CCR7 expression in EO771 cells did not affect cell growth under normal culture conditions *in vitro* (Fig. 1A) but EO771 CCR7 injected into the mammary fat pad displayed a dramatic increase in tumor growth *in vivo* compared to the control (Fig. 1B). Two major sources of tdTomato were observed in the TDLNs: tumor cells in the subcapsular sinus, which occasionally expanded into the LN parenchyma (Fig. 1C inset 1 and 3) distinguished by tdTomato expression and nuclear morphology, and accumulation of tdTomato as immune complexes associated with Follicular Dendritic Cells (FDCs) in B-cell follicles (Fig. 1C inset 2). In addition, tdTomato was detected in LN macrophages in both the vector control X2 and CCR7 positive tumors (Supplementary Fig. 1B). In primary tumors, lymphatic vessels were restricted to the tumor margins, where an association between tdTomato positive tumor cells and lymphatic vessels was observed (Fig. 1D).

**Fig. 1:**
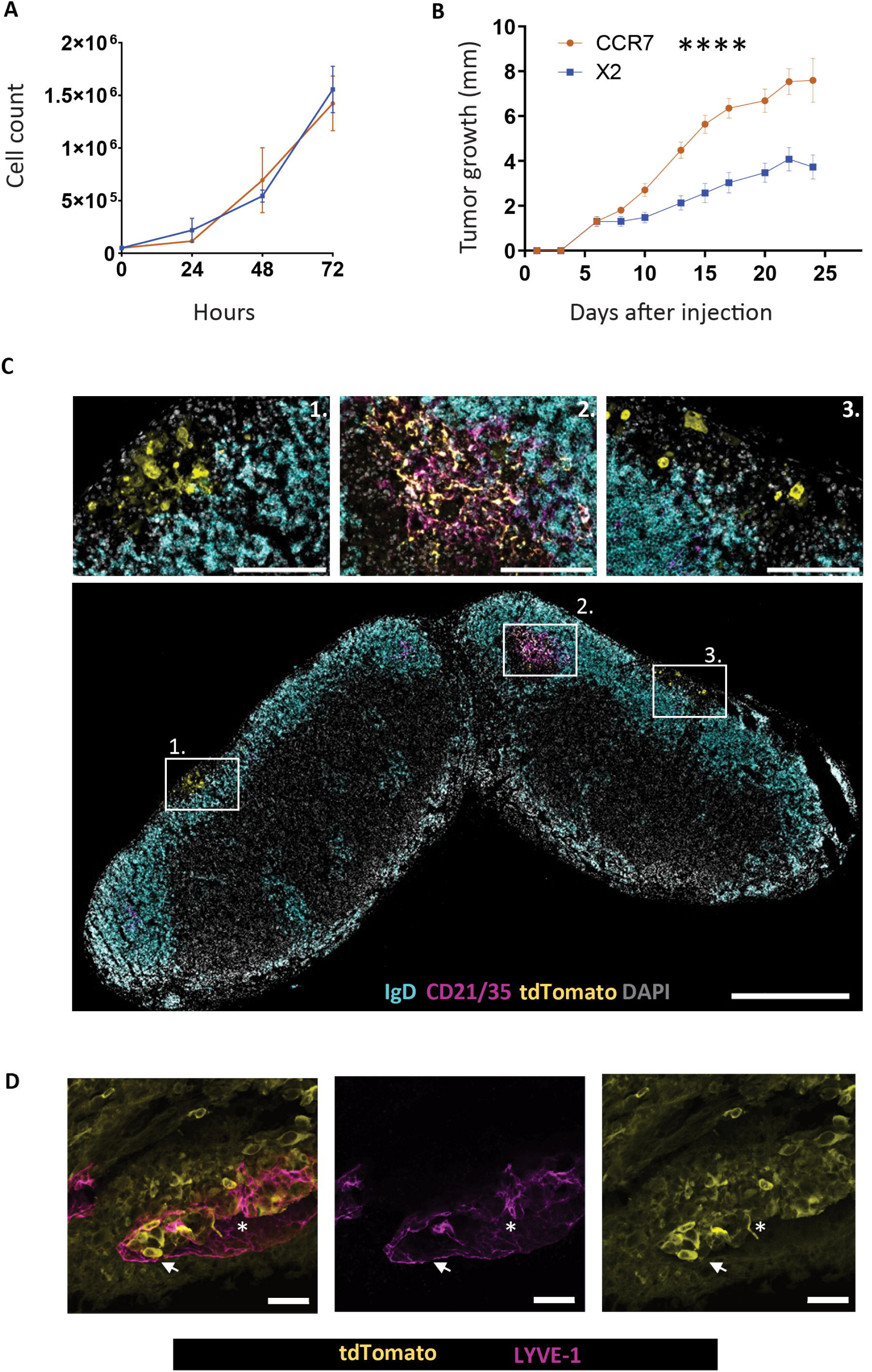
Expression of CCR7 in the triple-negative breast cancer cell line EO771 leads to increased tumor growth and lymphatic metastasis *in vivo*. A) *In vitro* proliferation of the X2 vector control- and the CCR7 expressing EO771. Data points indicate the mean values of three replicates and the error bars show the SD. Experiment representative of two Two-way ANOVA show no significant differences. B) Tumor growth *in vivo*. 15 mice per group were injected in the 4^th^ mammary fat pad with CCR7 or X2 control vector EO771. Tumor growth was monitored until the tumors reached >9 mm in diameter or day 24 after injection. Data points indicate the mean values with standard error of the mean (SEM). Non-parametric two-way ANOVA with Šídák’s multiple comparisons test show significant differences from day 10 and at all measured time-points up until day 24. P< 0.0001 **** C) tdTomato detected in LNs draining CCR7 positive tumors as metastatic cells (inset 1 and 3) and accumulation of tdTomato positive immune complexes on the surface of follicular dendritic cells (FDCs) in the B-cell zone (IgD) germinal centers (GCs) (inset 2). Scale bar overview 500 μm and insets 100μm. Representative of >4 LNs. D) Vibratome tumor section from CCR7 tumor stained for lymphatic vessels (LYVE-1) in magenta and tumor cells (tdTomato) in yellow. Scale bar 50 μm. Single plane optical section is shown. Arrow denotes tumor cells inside lymphatic vessels, star denotes vessel lumen. Representative of 3 stained tumors.

### Spontaneous upregulation of PDPN is needed for efficient lymphatic metastasis and enhanced *in vivo* growth of EO771 expressing CCR7

Staining for the glycoprotein podoplanin (PDPN), a marker of LN fibroblastic reticular cells (FRCs) and lymphatic vessels (24, 25), unexpectedly revealed that tumor cells reaching the subcapsular sinus (SCS) of TDLNs consistently expressed PDPN (Fig. 2A). Analysis of EO771 CCR7 cells by flow cytometry demonstrated the presence of a subpopulation of tumor cells with upregulation of PDPN, while the vector control only displayed scattered cells (Fig. 2B). In several tumor types, PDPN has been linked to increased invasion (24, 26). Although tumor cell-specific upregulation of PDPN has not been reported in human breast cancer *in vivo*, PDPN has been shown to be expressed by mammary epithelial basal cells, and loss of *Pdpn* was shown to reduce Wnt/beta-catenin-induced experimental tumors (27). To further explore the possible contribution of the PDPN phenotype to LN metastasis driven by CCR7, we sorted PDPN positive and negative cells from CCR7 expressing EO771 (Fig 2B). TdTomato expression levels were controlled for across the cell lines, as were CCR7 levels across PDPN-positive and PDPN-negative cell-derivates (Supplementary Fig. 2 A, B).

**Fig 2:**
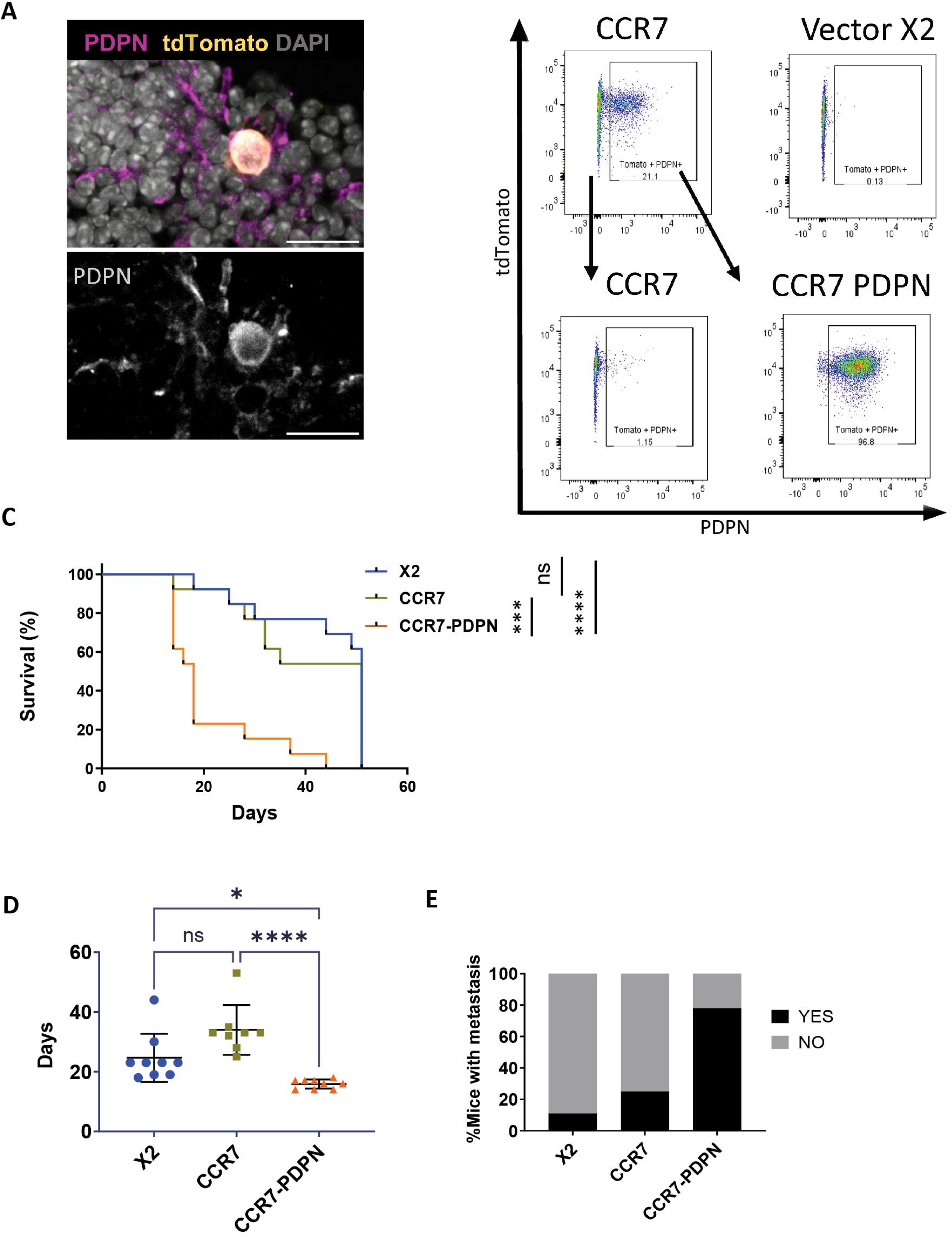
Spontaneous upregulation of PDPN is needed for efficient lymphatic metastasis and enhanced *in vivo* growth of EO771 expressing CCR7. A) Confocal image of PDPN (magenta) and tdTomato (yellow). Representative images of CCR7 expressing EO771 in TDLNs with expression of PDPN on arriving metastatic cells. Scale bar 20 μm. B) Cell sorting of PDPN positive and negative populations allowed the establishment of stable cell population of EO771 CCR7 with and without PDPN. Plots show representative FACS analysis for expression of tdTomato and PDPN. 21,1% EO771 CCR7, 0.13 % EO771 X2 vector control, 1.15% in EO771 separated CCR7 negative cells and 96,5% in EO771 CCR7-PDPN enriched for double positive cells. C) Survival curves. Mice were injected in the 4^th^ mammary fat pad. The tumors were allowed to grow until they reached a size of 9 mm. A log-rank Mantel-Cox test was used for statistical evaluation. n=13 in all groups. Significant differences were found between EO771 CCR7-PDPN and EO771 X2 (**** p <0.0001) as well as between EO771 CCR7-PDPN and EO771 CCR7 (*** p=0.0003). D) Time after tumor inoculation for samples used for determination of metastasis in E). Krusal Wallis test *p<0.05; ****p<0.0001. E) Metastasis determined by manual assessment of stained cryo-sections from at least 3 levels of each TDLN draining tumors within the same size range: i.e. i.e average 262 +/− 119 mg, X2 (n = 9), CCR7 (n = 8), CCR7-PDPN (n = 9) (62-100 sections per group).

Strikingly, survival based on time to reach terminal tumor size, was dramatically reduced in mice injected with CCR7-PDPN double-positive cells (18 days) compared to CCR7 or X2 vector control (both 51 days) (Fig. 2C). Tumor-take rates were also higher in CCR7 PDPN EO771 (92.3%) compared to CCR7 EO771 (61.5%) and vector X2 controls (38.4%). The latter was consistent across multiple experiments injections, with in average 50% for X2, 70% for CCR7 and 80% in CCR7-PDPN (Supplementary Fig. 2D). In contrast, no differences in tumor cell proliferation were observed in cell culture (Supplementary Fig. 2C).

To rule out the possibility that larger tumor size alone influenced metastasis, we compared metastasis with primary tumors selected for a medium size range (i.e average 262 +/− 119 mg across all groups), and by sectioning and imaging several levels of each TDLN. Imaging was chosen to differentiate tdTomato-positive immunocomplexes on FDCs and tdTomato-positive macrophages from real metastasis with high confidence. Even with a significantly shorter time for tumor development (Fig. 2C), CCR7-PDPN tumors displayed a higher incidence of metastatic spread to TDLNs than CCR7 and X2 controls (Fig. 2E). In contrast, the accumulation of immunocomplex-associated tdTomato on FDCs, LN size or GC formation did not differ across the groups (Supplementary Fig. 3A-E). The latter was also the case when tumors were harvested at the same timepoint (day 14) (Supplementary Fig. 3F-J). The accumulation of tdTomato across the GC or FDC area (an indicator of ongoing GC formation and B-cell activation (28)) was the same across groups despite differences in tumor size (Supplementary Fig. 3F-J). This indicate that lymphatic transport and accumulation of immune complexes on FDCs are independent of tumor size and metastatic spread to the TDLNs. Taken together, these data support that CCR7 expression in EO771 increases metastatic spread to the TDLNs, but this effect requires spontaneous upregulation of PDPN.

### Spontaneous PDPN upregulation in EO771 does not drive tumor growth or lymphatic metastasis without CCR7

To exclude the possibility that PDPN upregulation alone would be sufficient to drive metastasis, we established PDPN-positive vector control cells. In the X2 vector control, despite no defined population of PDPN-expressing cells, sparse cells were consistently stained (Fig. 2B and Fig. 3A). By subsequent sorting for PDPN positive cells, an enrichment was obtained that remained stable in culture, varying between ∼80-90% (Fig. 3A and data not shown). PDPN negative and positive EO771 showed no difference in tumor growth *in vivo* (Supplementary Fig. 4A and Fig. 3B-3C), but did display a higher tumor-take based on detectable palpable tumors: 5/14 injections i.e. 35.7% X2, versus 11/14 injections i.e. 78,5 % for X2 PDPN, at day 14 after tumor cell injection. However, the increased growth seen in EO771 cells co-expressing CCR7 and PDPN could not be replicated by PDPN alone (Fig. 3C). Similar to the original vector control, X2 PDPN tumors, required a significantly longer time to develop to end size, compared to CCR7-PDPN, and did not display LN metastasis (Supplementary Fig. 4B, C).

**Fig. 3:**
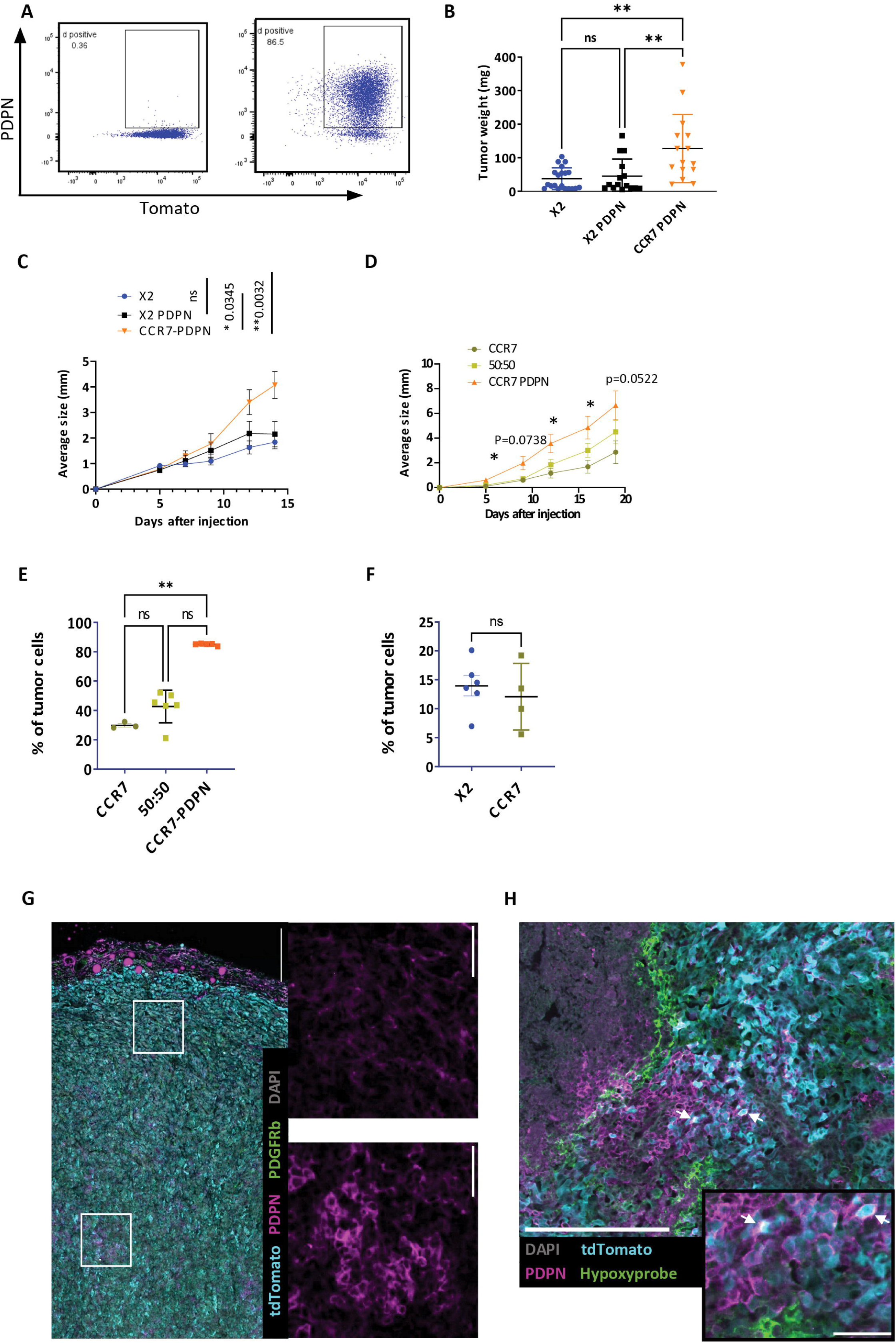
Spontaneous PDPN upregulation in EO771 does not drive tumor growth without CCR7. A) Original proportion of PDPN+ cells in EO771 vector control X2 0.36% (left) and increased proportion established by FACS sorting 86.5% (right). B-C) Mice were injected with EO771 cell line derivates: X2 (n=12), X2 PDPN (n=8), and CCR7-PDPN (n=8), on both sides in the 4^th^ mammary fat pad. Tumor growth was monitored up until day 14. B) Tumor weight at end stage for the different groups. n=20 X2, n=15 X2 PDPN, n=15 CCR7-PDPN. Kruskal Wallis test and Dunn multiple comparison were used for statistical analysis. ** p< 0.01. C) Growth curve. Average diameter is shown. Data points show the mean, and error bars the standard deviation (SD). A non-parametric Mixed effect analysis was used for statistical analysis and p-values for day 14 are shown. D) Injection of 3 tumor groups: CCR7, CCR7-PDPN and a 50:50 mix of CCR7 and CCR7-PDPN EO771, n=5 mice in each group with injections on both sides in the 4^th^ mammary fat pad. Significant differences in tumor growth is seen between CCR7 and CCR7-PDPN while the 50:50 group show intermediate growth. E) Proportion of PDPN positive tumor cells in end stage tumors. Kruskal Wallis test and Dunn multiple comparison were used for statistical analysis. ** p< 0.01. F) Induction of PDPN in size matched tumors from mice injected with PDPN negative X2 vector control or CCR7 positive EO771. No significant differences. G) Staining of EO771 X2 tumor for tdTomato (cyan), PDPN (Magenta), PDGFRbeta (green) and Nuclei (DAPI grey). Scale bar 200 μm. Insets show central foci (1) with PDPN expression and peripheral foci (2) with low or no detection. Scale bar 50 μm. H) Staining of a EO771-CCR7 tumor for hypoxi-probe (green) together with detection of tTomato (cyan), PDPN (magenta) Nuclei (DAPI grey). Hypoxi-probe positive cells (arrow) line the necrotic areas (star). PDPN is seen expressed in cells adjacent to these areas, and in rare hypoxic cells (arrowhead). Scale bar 200 μm, inset 50 μm.

To evaluate if CCR7 expressing cells with PDPN have a direct growth advantage *in vivo,* we conducted a competitive experiment by injecting mixed cells, 50:50 CCR7:CCR7-PDPN, with CCR7-PDPN alone or CCR7 alone as controls, and evaluated the proportion of PDPN expression in end-stage tumors. Surprisingly, no selection was evident, which suggests that there was no direct growth or survival advantage for double positive cells (Fig. 3D and E). However, PDPN positive cells were unexpectedly observed also in tumors derived from the injection of PDPN-negative CCR7 tumor cells (Fig. 3E). Intriguingly, this induction was still not sufficient to drive increased growth to the same extent as when CCR7-PDPN-positive cells were injected (Fig. 3D).

The spontaneous induction of PDPN *in vivo* was unexpected. We confirmed that the induction of PDPN also occurred in the X2 vector control (Fig. 3F), with individual tumors showing larger variation than the groups (Fig. 3E and 3F). *In situ* evaluation indicate that PDPN-positive tumor cells were primarily found in the central parts of the tumor mass in groups of cells, both in CCR7 and X2 vector control tumors (Fig. 3G and data not shown).

PDPN has been previously suggested to be linked to mammary stem cell functions (27). However, we did not detect any difference between the four cell derivatives in colony forming assays *in vitro* (Supplementary Fig. 5). Intra-tumoral induction of PDPN *in vivo* may indicate a link to hypoxia. To explore this possibility, mice were injected with Hypoxyprobe (Fig. 3H and Supplementary Fig. 6).

Hypoxia is expected to be most pronounced in the center of larger tumors, which was also evident by the common absence of live (tdTomato-positive) tumor cells in the center of EO771 tumors (Supplementary Fig. 6). Hypoxyprobe-positive cells were detected along the border between live and necrotic tumor areas (Fig. 3H and Supplementary Fig. 6). While PDPN expression was only rarely observed in hypoxic cells, it was consistently detected in clusters of cells adjacent to these regions. Consistently, tumor cells at the tumor rim, outlined by PDPN-positive fibroblasts, remained negative or low in PDPN. Together, these data suggest an interplay between hypoxia and tumor plasticity within the tumor microenvironment of EO771.

### CCR7 and PDPN double positive tumors display a shift to a more immune-cold profile

CCR7-PDPN cells were not selected for in competitive experiments with CCR7 single-positive cells (Fig. 3D and E), indicating that other mechanisms allow increased tumor progression *in vivo*. The proportion of tumor infiltrating lymphocytes (TILs) in human primary tumors has been used to define so-called immune-cold (low in TILs) versus immune-hot (high in TILs), where immune-hot tumors in general respond better to immunotherapy (29). This reflects the need for tumor cell-lymphocyte interactions for cellular tumor immunity. EO771 forms a capsule with a high density of fibroblast, myeloid cell and T-cells, while the density of TILs is sparse (Fig. 4A and data not shown). The latter is particularly evident in early stage tumors. While flow cytometry loses the spatial context, we chose to analyze CD8 TIL density based on imaging. Analysis of tumors 14 days after injection showed reduced high-density areas of intra-tumoral CD8 T-cells in CCR7-PDPN compared to the vector control, as assessed by analysis of random areas across the tumor excluding the tumor border (capsule) and central necrotic areas (Fig. 4C, D). This difference persisted when comparing end stage tumors, suggesting that it is inherent to the tumor type rather than reflection of tumor size (Fig. 4E, F). Hence CCR7-PDPN tumors differentiate from vector control by reduced high-density areas of intra-tumoral CD8 T-cells, suggesting a shift to a more immune cold profile. In contrast to CD8 T-cells, CD11b (macrophages), which constitutes the major population of immune cells in EO771, did not show any clear reduction in CCR7-PDPN tumors compared to the control (Supplementary Fig. 7A-B). In addition, vascular density in CCR7-PDPN compared to the vector control, as determined by flow cytometry, showed no differences (Supplementary Fig. 7C).

**Fig. 4:**
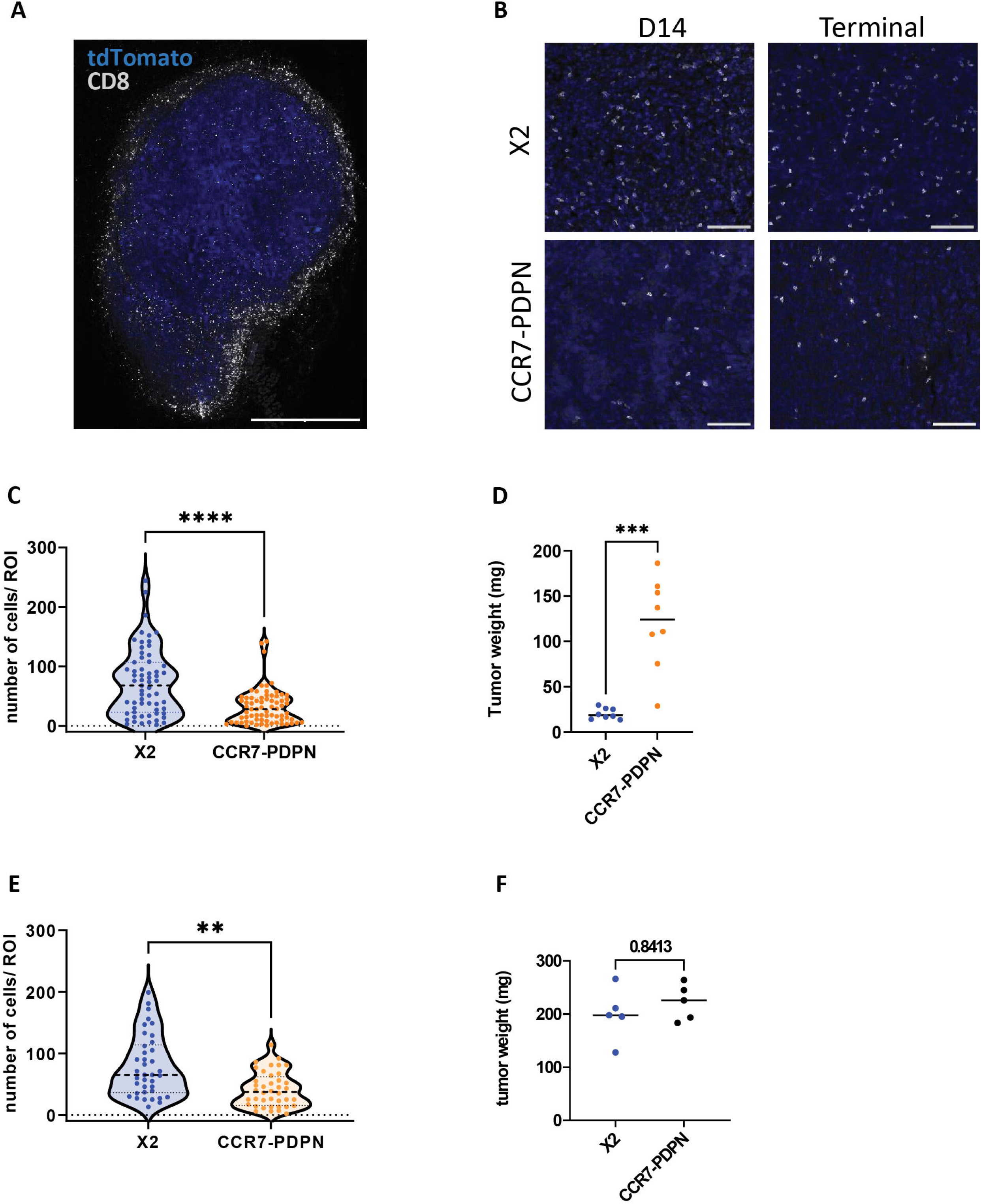
CCR7 and PDPN double positive tumors display a shift to a more immune-cold profile. A) Overview of a tumor at an early stage, stained for CD8 (white) and tdTomato shown in blue visualizes also the tumor-cell free border with high density of T-cells. Scale bar 1 mm. B) Representative images of CD8 staining (white) and DAPI (nuclei) in blue, central tumor areas. Scale bar 100 μm. C) Quantification of T-cell density day 14 after tumor injection, 63 and 72 areas per group, n=8 each group. D) Tumor weight of analyzed samples in C) n=8 each group. E) Quantification of T-cell density in size matched tumors, 38 and 40 areas (Regions of Interest ROIs) per group. N=5 each group F) Tumor weight of analyzed samples in E, n=5. C-F Krusal-Wallis test. **** p<0.0001, *** p<0.001, ** p<0.01

### CCR7 and PDPN Expression in EO771 Cells: Impacts on EMT Signature, Collagen Expression, and the IFN Response

PDPN could either be upregulated alone or be part of a differentiation program with other molecular changes in EO771 cells. To distinguish between these possibilities and further understand the molecular consequences of CCR7 expression and spontaneous PDPN upregulation in EO771 cells, we performed comprehensive RNA profiling.

Principal component analysis (PCA) revealed distinct clustering of the four cell derivatives with and without CCR7 and PDPN (Fig. 5A). The tumor classification of EO771 mammary carcinoma has been debated, but it has most often been assigned as a TN breast cancer (TNBC) or basal cell-like phenotype (18, 19). Our data revealed very low, near-background mRNA levels of Estrogen Receptor (ER) alpha (*Esr1*) and no detectable levels of ER-beta (*Esr2*). Progesterone receptor membrane components (*Pgrmc1, Pgrmc2*), HER2 (*Erbb2*) and HER3 (*Erbb3*) were all expressed at variable but not high levels (Supplementary Fig. 8). Notably, EO771 cells exhibited robust expression of several EMT-associated markers (22), including *Vim, Twist1, Zeb1/2, and Wnt7b* (Supplementary Fig. 8). In contrast, cytokeratins were expressed at very low or undetectable mRNA levels, except for Keratin 8 (*Krt8*) (Supplementary Fig. 8). Notably, although EO771 exhibited very low mRNA levels of ERα (*Esr1*), this expression was further reduced in CCR7-PDPN double-positive cells (Supplementary Fig. 8). A similar pattern was observed for *Erbb2*, suggesting further suppression of mammary receptor expression in the context of CCR7 expression.

**Fig. 5:**
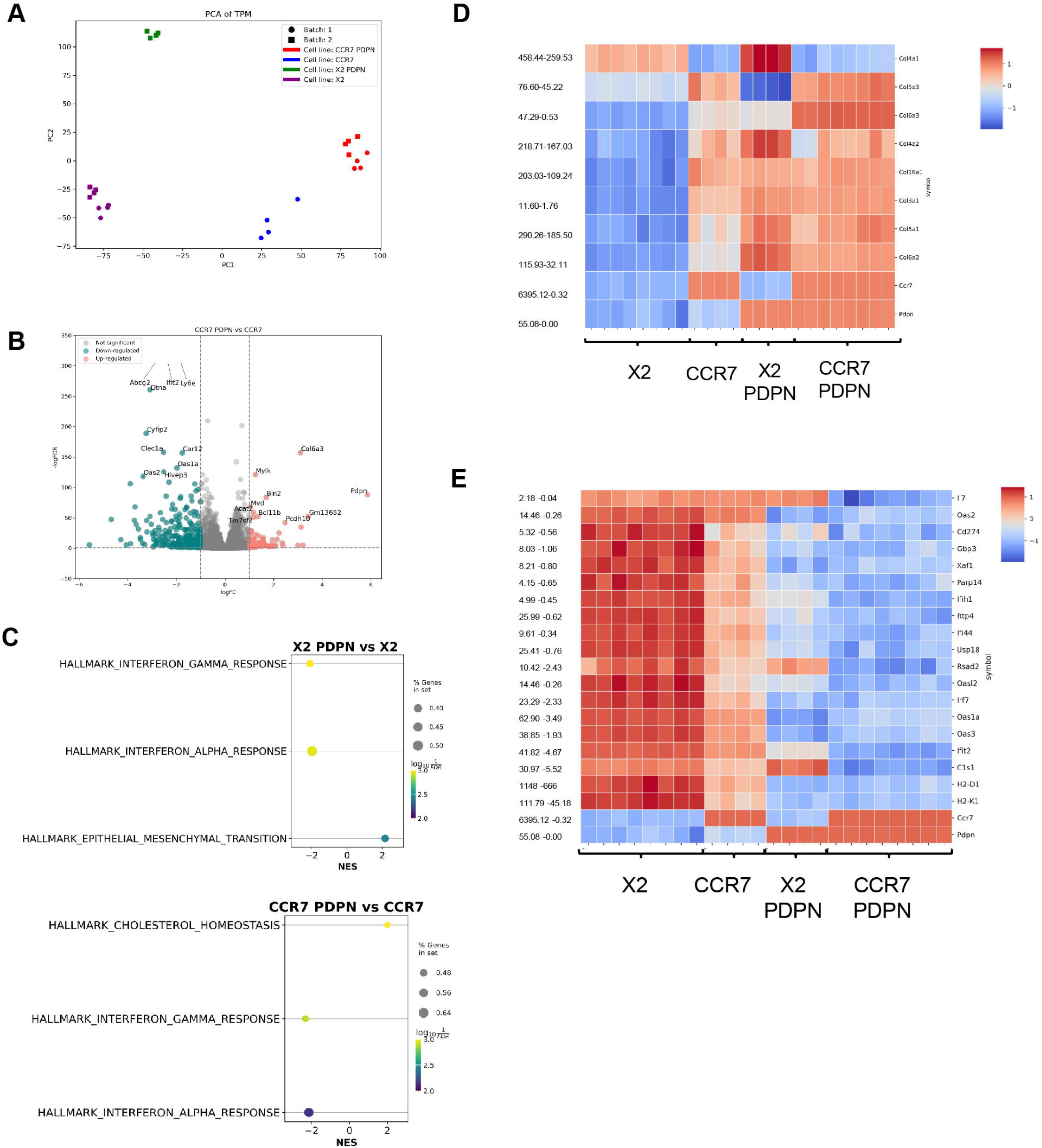
CCR7 and PDPN Expression in EO771 Cells: Impacts on EMT Signature, Collagen Expression, and the IFN Response. A) Principal Component Analysis (PCA) shows the distribution of four EO771 cell lines across two experimental batches. Data points are colored and grouped by cell type and batch. B) Volcano plots displaying differentially expressed genes (DEGs) between CCR7-positive and CCR7-PDPN positive EO771 cells. Dashed lines indicate thresholds for upregulation and downregulation (∣Log2FC∣>1) and statistical significance (adjusted p-value < 0.05). C) Gene Set Enrichment Analysis (GSEA) using mouse hallmark gene sets highlights differences between PDPN-positive and PDPN-negative cell lines. D-E) Heatmaps displaying Z-scores of normalized gene expression, where Z-score = 0 represents the mean gene expression. Gene names are indicated on the right side, normalized gene expression values are added on the left side of the heat maps for reference. EMT-related collagens in CCR7 and PDPN positive EO771 cell derivatives (D). IFN-related and immune-related genes (i.e., *Il-7*) (E).

Gene Set Enrichment Analysis (GSEA) using mouse hallmark gene sets showed an enhanced EMT signature in the PDPN-positive vector line (X2 PDPN) compared to the original X2 control (Fig. 5C). The enhanced EMT in PDPN-expressing EO771 cells indicates that PDPN is upregulated as part of a shift in cell differentiation. In line with this, the top significantly upregulated differentially expressed genes (DEGs) in EO771 with PDPN expression encoded for collagens (Supplementary Table 1). The gene expression profile of collagens was further modulated by the co-expression of PDPN and CCR7, as illustrated by the changes of selected EMT-associated collagens (Fig. 5D).

Among the top-ranked hallmark pathways FDR <0.05), only cholesterol homeostasis showed an increase in CCR7-PDPN positive EO771 cells (Fig. 5C). The top-ranked downregulated hallmark pathways included both type I and type II interferon (IFN) responses (Fig. 5C). Downregulation of IFN response-associated genes was evident in cells expressing either PDPN or CCR7 alone compared to the vector control and was further reduced in cells co-expressing both markers (Fig. 5E). CCR7-PDPN cells also showed a downregulation of *Il7* expression (Fig. 5E). The observed immune-modulation extended to MHC class I molecules, with reduced expression of H2-D1 and H2-K1 in CCR7-PDPN cells.

In conclusion, CCR7 expressing EO771 cells with spontaneous PDPN upregulation exhibit features suggesting further dedifferentiation, altered EMT-associated collagen expression, and an immunomodulatory profile with the suppression of baseline IFN signaling. An overview of the top 100 up-regulated and down-regulated, DEGs across the four cell lines is provided in Supplementary Table 1.

### Heterogeneous expression of PDPN is linked to TN breast cancer and EMT in Human Breast Cancer

Although CCR7 has been extensively studied as a factor that promotes lymphatic metastatic behavior in human breast cancer (8, 9, 11, 12), PDPN has not been thoroughly studied. Therefore, we performed *in-silico* analyses to explore the potential translational relevance of PDPN expression in breast cancer cells.

We first explored a single cell RNAseq data set of 32 human breast cancer cell lines (Supplementary Fig. 9A) (30). Although several of the cell lines in this data set have been reported to express CCR7, i.e. MCF-7, DU-4475 and MDA-MB-361 (8, 12), CCR7 was not detectable at the single cell mRNA level (data not shown and (30)). PDPN was detected in two TNBC cell lines, CAL-51 and MDA-MB-436, and in the basal-like pre-malignant cell line MCF-12A (Supplementary Fig. 9B), which is consistent with the known expression of PDPN in mouse mammary basal epithelial cells (27). Both CAL-51 and MDA-MB-436 were originally isolated from metastatic sites (30). Although sub-clustering did not separate PDPN-positive from PDPN-negative cells, the data supports heterogeneous expression of PDPN in human TNBC cell lines (Supplementary Fig. 9C).

We next used a single cell data set of neoadjuvant treated ER+, HER2+ and TNBC from 26 patients (31). CCR7 was not detectable except in immune cells. However, PDPN-expression in tumor cells could be confirmed in one patient with a metaplastic subtype of TNBC (Fig. 6A). To ensure that subsequent analyses were not confounded by individual or treatment specific differences between patients we focused on performing intra-patient analysis of this patient. Unsupervised Leiden gave the separation of the cancer cells into two clusters. PDPN cells were scattered in both clusters but were significantly enriched in cluster 0 (∣Log2FC∣>1 and adjusted P-value < 0.05|) (Fig. 6A and 6B). GSEA analysis using human hallmark gene sets showed upregulated hallmarks for Transforming Growth Factor beta (TGFβ), EMT and hypoxia (Fig. 6C). Collagen analysis revealed a similar pattern of upregulation as in EO771 with PDPN expression (Fig. 6D). Our data support that human breast cancer can display heterogenous upregulation of PDPN which mirrors the effects we described in PDPN-positive EO771, including changes in collagen expression and shifts in the EMT profile.

**Fig. 6:**
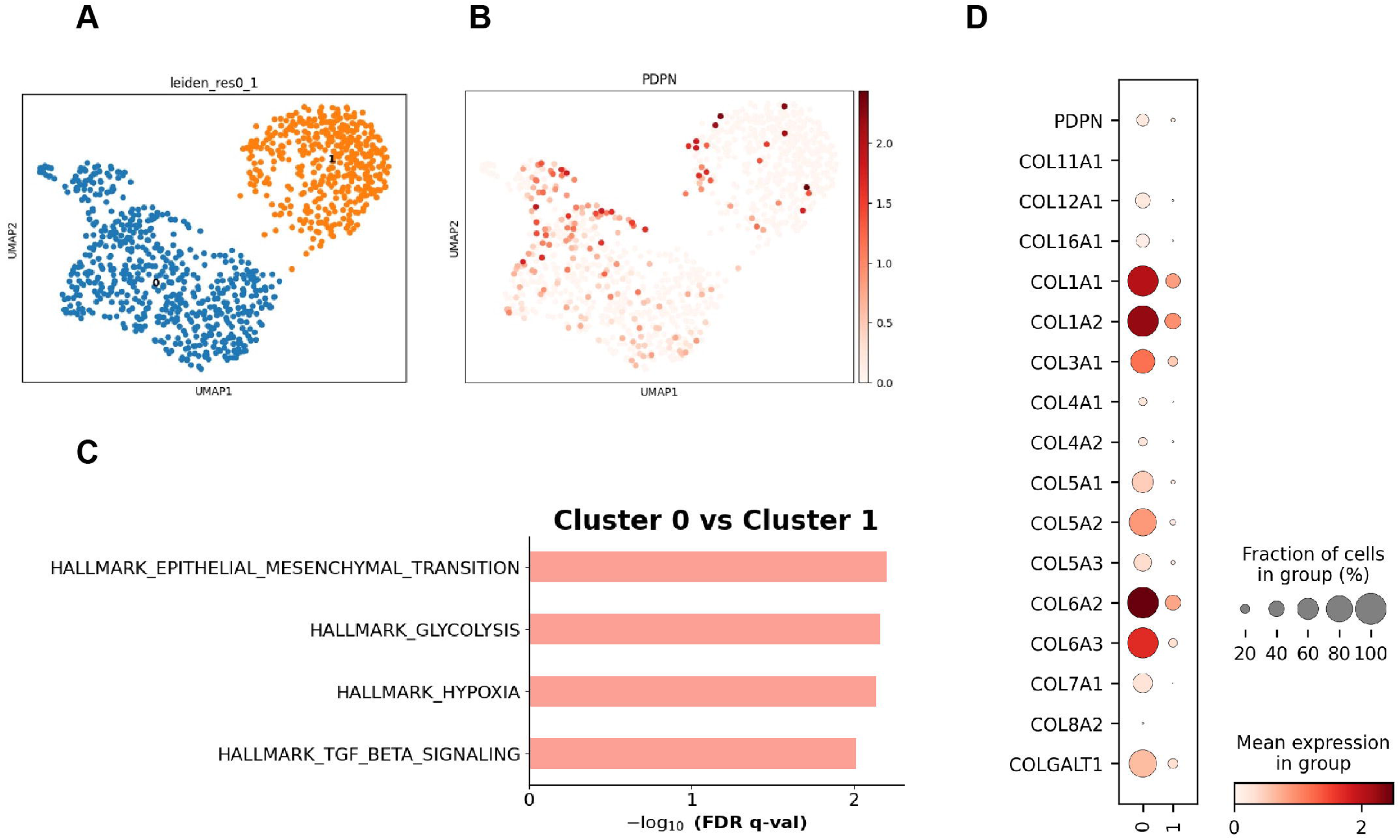
Single-cell transcriptional analysis of metaplastic triple-negative breast cancer (TNBC) reveals heterogeneity in PDPN expression that is linked to shifts in EMT. A) UMAP visualization of single-cell transcriptomic data segregates cells into two distinct clusters based on Leiden clustering (resolution = 0.1) (Cluster 0, blue; Cluster 1, orange). B) UMAP plot showing the expression of PDPN across the dataset, with higher expression levels concentrated in Cluster 1. C) Hallmark pathway enrichment analysis comparing Cluster 0 and Cluster 1. Significant pathways upregulated in Cluster 0 include epithelial-mesenchymal transition (EMT), glycolysis, hypoxia, and TGF-β signaling. D) Dot plot showing the expression of PDPN and selected collagen genes in the two clusters. Dot size indicates the fraction of cells expressing the gene within each group, and color intensity represents mean expression level.

### *In silico* analyses show that high CCR7-PDPN mRNA expression score associates with aggressive tumor characteristics and presents independent prognostic value

Given synergistic effects between CCR7 and PDPN in the EO771 model for both tumor growth and LN metastasis, we further evaluated their cooperation in human breast cancer progression.

To this end, we explored their joint prognostic potential using the Molecular Taxonomy of Breast Cancer International Consortium (METABRIC) datasets (32) with two groups: discovery (n=939) and validation (n=845). We established a signature score featuring the summarized expression values of CCR7 and PDPN, creating a CCR7-PDPN mRNA expression score (CCR7-PDPN score). The METABRIC cohorts are based on expression data derived from tumor cores, which may contain varying proportions of stromal and immune cells, although they are selected for areas of high tumor cell content. As a result, both stromal and tumor cells may contribute to the observed correlations. Therefore, the analyses performed on the METABRIC cohorts should be regarded as exploratory.

To evaluate how the high CCR7-PDPN score (cut-point upper quartile) was related to tumor phenotypes, common associations in both discovery and validation groups included, high histologic grade (p<0.001 Fig. 7 A-B), ER negativity (p=0.001 Fig. 7C-D), and aggressive molecular phenotypes (p=0.001 Fig. 7G-H). Additionally, we demonstrated associations with LN metastases (METABRIC discovery cohort, p=0.009 Figure 7G), and low tumor size (METABRIC validation cohort, p=0.031 Fig. 7H). High CCR7-PDPN expression score also associated with shorter disease-specific survival (univariate analysis, METABRIC validation cohort, Fig. 7I, Supplementary Table 2A).

**Fig. 7:**
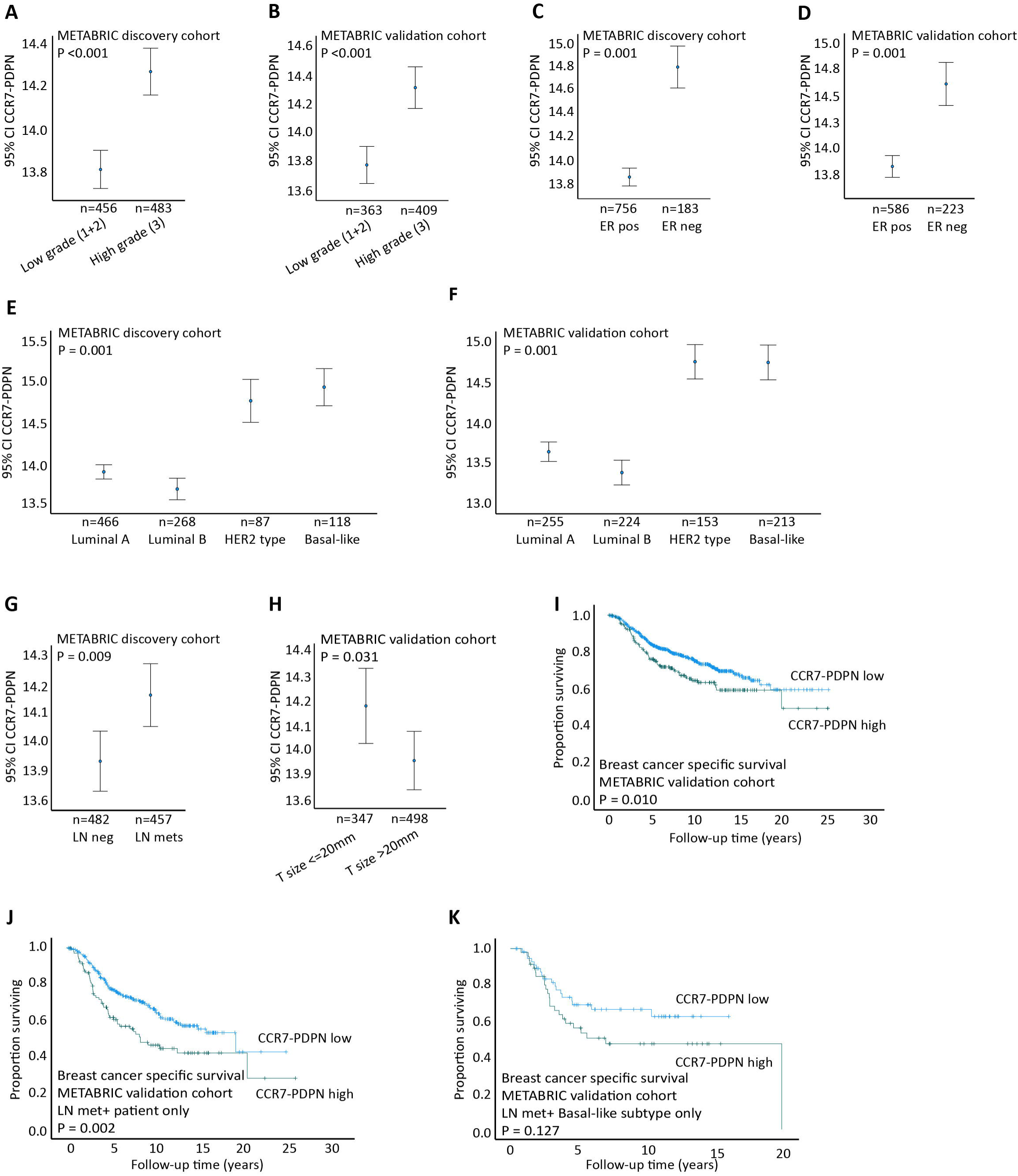
CCR7/PDPN across clinico-pathological variables, and breast cancer specific survival according to CCR7/PDPN levels. A-F) CCR7/PDPN score across histologic grade (A-B), ER status (C-D), and molecular subtypes (E-F) in METABRIC discovery and validation cohorts. G-H) CCR7/PDPN score across lymph node status (G) and tumor size (H) in METABRIC discovery and validation cohort, respectively. I) A high CCR7/PDPN score associated with shorter disease-specific survival (METABRIC validation cohort). J) A high CCR7/PDPN score was also significantly associated with shorter survival in lymph node positive tumors only. K) A trend of significance was observed in lymph node positive, basal-like subtype patients. All Kaplan-Meier survival plots from the METABRIC validation cohort. Data shown with error bars representing 95% confidence interval of the mean, and p-values by Mann–Whitney U test.

Interestingly, when adding the clinico-pathological variables tumor diameter, histologic grade, ER status and lymph node status to the multivariate analysis, a high CCR7-PDPN score was demonstrated independently associated with shorter disease-specific survival (METABRIC validation cohort; HRC=C1.40, 95% CI 1.03-1.91, pC<C0.029, Supplementary Table 2A). Moreover, when investigating the prognostic impact of CCR7-PDPN score in LN status met-/met+ separately, our data showed that high CCR7-PDPN score was associated with reduced survival in LN metastasis positive patients (METABRIC validation cohort; HR = 1.81, 95% CI 1.28-2.58, p=<0.001, Supplementary table 2B, Fig. 7B), but not in LN negative patients (data not shown). When investigating the prognostic impact of CCR7-PDPN in individual molecular subtypes, high expression of CCR7-PDPN score demonstrated a trending significant association with shorter disease-specific survival in LN positive, basal-like subtype (METABRIC validation cohort; Fig. 7K), but not in the other subsets.

## Discussion

EO771 triple negative mammary carcinoma, like most models of cancer in a C57Bl6 background has limited lymphatic metastatic potential (18–20). We showed that CCR7 expression conditions the cells for enhanced *in vivo* tumor growth and LN metastasis. However, this effect was only effective in CCR7-positive EO771 that also displayed spontaneous upregulation of the glycoprotein PDPN. Neither CCR7 expression nor PDPN upregulation were sufficient alone, underscoring a cooperative role in enabling efficient metastatic spread.

CCR7 has been recognized as a contributor to lymphatic metastasis for over two decades (8), however its reliability as a biomarker remains inconsistent (17). Despite the high EMT features of EO771, including the expression of vimentin and low levels of keratins, our data showed that CCR7 expression alone is not sufficient to drive tumor progression and LN metastasis. Instead, a further shift in the mesenchymal phenotype, marked by PDPN upregulation, is required. EMP, characterized by reversible shifts between epithelial and mesenchymal states (21, 22), which we demonstrated to occur spontaneously *in vivo* in the hypoxic TME, appears to be a crucial factor in CCR7-driven metastasis in this context. In both mouse and human TNBC, we showed that PDPN expression correlates with increased EMT scores and collagen expression, reinforcing the translational relevance of these mechanisms. We also showed that these effects were further modulated by the co-expression of PDPN and CCR7 in EO771 cells.

PDPN primarily interacts with CLEC-2-expressing immune cells and platelets. In experimental melanoma, PDPN expression has been linked to increased tumor invasiveness, cellular migration and/or mobility (26) and platelet interaction in hematogenous metastasis (33). It has also been suggested to be a stem-like tumor cell marker (27). However, under normal growth conditions we did not find any changes in growth, cell morphology or colony formation assays in the EO771 model. Specific changes as part of the EMT shift and PDPN expression, or a combination of changes, that are critical for the observed effects on growth and metastasis in CCR7-PDPN co-expressing EO771 *in vivo*, remain to be defined in future studies. One contributing factor may be the modulation of tumor-expressed collagens.

Although collagens are mainly associated with cancer-associated fibroblasts (CAFs), their upregulation in tumor cells has been linked both to proliferation and metastatic niche formation (34, 35). Interestingly EO771 cells arriving at the TDLNs were consistently PDPN-positive, even when the primary tumor consisted of mixed cells. In EO771 with co-expression of CCR7 and PDPN, the interplay between ECM components and tumor plasticity likely contribute to establishing a microenvironment conductive to metastasis. Recent data from 4T1, a TN model in a Balb/c genetic background, support that early metastatic spread is driven by tumor cells with high EMT features while the outgrowth of the metastatic lesion inside the LN can favor a switch to a more epithelial phenotype (36). Future follow-up studies on these questions and dynamic changes will be of interest across different tumor models, including EO771 CCR7-PDPN, and human cancer.

Our analysis revealed suppression of type I and II interferon responses, suggesting an immune evasion phenotype of EO771 with expression of PDPN and CCR7. Supporting this notion, tumors co-expressing CCR7 and PDPN exhibited an immune-cold phenotype characterized by reduced infiltration of tumor-infiltrating lymphocytes (TILs). That cell-intrinsic restriction of IFN responses can have an inhibitory effect on the anti-tumor response has also been demonstrated in human cancer (37). These responses can be triggered by cytosolic nucleic DNA in cancer cells with high genomic instability through damage-associated signaling, such as the cGAS/STING system (38) which has also been associated with tumor dormancy and reactivation (39). Knockdown of PARP7 (inhibitor of IFN pathways) in EO771 was recently shown to drive increased tumor immunity *in vivo*, which also support that this tumor model is dependent on immune evasion via IFN pathway suppression (40). The accumulation of immune complexes and B-cell GC formation, previously linked to the promotion of tumor growth in this EO771(41), was unchanged in tumors formed with and without CCR7 and PDPN. This indicate that modulation of B-cell activation does not explain the observed differences, and also suppport that general lymphatic transport to the TDLNs is unchanged across EO771 derivatives with and without CCR7 and PDPN.

The role of tumor-expressed PDPN in the progression of different types of human carcinomas has been suggested in multiple studies (24), but no studies on primary human breast cancer have been published to date. Corroborating the translational relevance of our findings, heterogeneous PDPN expression was also detected in human breast cancer cell lines, specifically TN and basal-like cells, and one example of primary breast cancer. The latter is a TN breast cancer of the metaplastic subtype, which is a rare but highly aggressive form of breast cancer, with a mixed histological phenotype from epithelial to spindle and mesenchymal (42). LN metastasis is not as common in TN breast cancer, including metaplastic breast cancer, as it is in luminal breast cancer. However, LN metastasis is also associated with worse prognosis in this tumor type, and increased EMT has been linked to nodal metastasis (43). In mice, mammary basal epithelial cells were shown to express PDPN, and epithelial PDPN was also linked to promotion of tumorigenesis (27). In human few studies have been performed, but non-tumorigenic human basal cell lines have been shown to display cellular plasticity *in vitro* with differentiation into mesenchymal-like cells through EMT with upregulation of PDPN (44). Hence, basal-like and TN metaplastic breast cancers are candidate subtypes for future studies.

Supporting this notion, our results showed an association between the CCR7-PDPN mRNA score and the basal-like subtype in the METABRIC analyses. We showed that high CCR7-PDPN scores correlated with aggressive tumor features, including TNBC, and to reduced survival in LN-positive patients. A high CCR7-PDPN score also demonstrated an independent prognostic impact in multivariate analysis, when stratifying for LN status. Notably, CCR7 and PDPN as individual markers were not significant in the same tests. To our knowledge, this is the first study to describe the expression and prognostic value of CCR7-PDPN in breast cancer patients paired with well-characterized clinico-pathological variables and a long and complete follow-up information. Although the association between the METABRIC CCR7-PDPN score and poor prognosis demonstrated in our study, could also be partly dependent on CAFs which are a major source of PDPN in the tumor microenvironment together with lymphatic vessels, a contribution from PDPN-expressing tumor cells is also possible. These data motivate and form a basis for follow-up studies with a detailed analysis of the cell types contributing to the observed association.

Taken together, by linking tumor microenvironment-induced shifts in EMT and PDPN expression to metastatic potential, we provide new insights into new molecular determinants of interest for breast cancer progression and demonstrate the translational relevance for human breast cancer. These results underscore the importance of tumor plasticity in shaping CCR7 chemokine receptor-driven lymphatic metastasis and immune evasion.

## Materials & Methods

### Cell Culture and Cell Sorting

The EO771 CCR7 cell line and its empty vector control, EO772 X2, were generated through retroviral transduction. Specifically, PLNCX2 (Clontech) vectors containing either mouse *Ccr7* cDNA or the tdTomato reporter gene were transduced into EO771 mouse mammary carcinoma cells (CH3 Biosystems). Retrovirus production was performed using Phoenix ampho-helper-free cells (developed by Garry P. Nolan, Stanford University). Cells expressing tdTomato and CCR7 were enriched by fluorescence-activated cell sorting (FACS) on a BD FACSAria III (BD Biosciences) with a 100 µm nozzle and 20 psi pressure setting. CCR7 was detected using anti-CCR7-APC (clone 4B12, Thermo Fisher Scientific) and dead cells were excluded using SYTOX™ Blue dead cell stain (Thermo Fisher Scientific). Further sorting was conducted to isolate cell populations based on spontaneous upregulation of PDPN. Cells were stained with anti-PDPN-eFluor™660 (clone eBio8.1.1) to identify PDPN-positive populations in CCR7 or X2 lines, yielding four distinct cell lines: X2, X2 PDPN, CCR7 and CCR7-PDPN. No clonal selection was performed. Instead, bulk sorting was used to retain the overall heterogeneity of each population. Over the course of repeated experiments both CCR7 expression and PDPN expression were controlled by flow cytometry.

Cells were maintained in RPMI-1640 with HEPES (Gibco™), supplemented with 10% fetal bovine serum (FBS, Gibco™), 1% L-glutamine (Gibco™), and 1% penicillin-streptomycin (Gibco™) at 37°C in a humidified incubator with 5% COC. Cells carrying the vector constructs were cultured under selective pressure with 1 mg/ml G418 (Invitrogen, Thermo Fisher Scientific). Proliferation was measured manually by seeding a total number of 0.5×10^5^ cells per well in triplicates in a 12-well plate. Cells were harvested at 24-, 48- and 72-hours cells were harvested. Live cells were counted using Countess II Automated Cell Counter (Invitrogen). Alternatively, the CyQUANT® Cell Proliferation Assay Kit (Invitrogen) was used according to the manufacturer’s instructions.

### RNA seq and bioinformatical analysis cell lines

RNA sequencing was conducted at the Bioinformatics and Expression Analysis (BEA) core facility at the Karolinska Institute. Total RNA was extracted from quadruplicate EO771 samples (technical replicates), using the RNeasy Mini Kit (Qiagen, Valencia, CA) following the manufacturer’s instructions. Libraries were generated using the TruSeq Stranded mRNA kit (Illumina) and sequenced on an Illumina NextSeq 550 platform in single-end mode, producing 75 bp reads mapped to the mouse reference genome (GRCm38/mm10).

Four cell lines were divided into two batches for sequencing: the first batch included X2, CCR7, CCR7-PDPN and the second batch included X2, X2 PDPN, and CCR7-PDPN. All the data was analyzed using Python (v.3.11.5). Normalization was performed using Transcripts Per Million (TPM) by rnanorm (v.1.5.1). TPM values below 1 were considered as noise (25). Subsequently, principal component analysis (PCA) was conducted using sklearn (v.1.3.0) to reduce dimensionality and visualize the data, particularly by observing batch effects and sample variations. Differentially expressed genes (DEGs) analysis was performed using pyDESeq2 (v.0.4.4). DEGs exhibiting an absolute log2 fold change >1 (|log2FC| > 1) and adjusted p-value < 0.05 or false discovery rate (FDR) < 0.05 were regarded as significant. Gene set enrichment analysis (GSEA) was performed using the GSEApy (v.1.1.1), with mouse hallmark gene sets (v2023.2) obtained from the Molecular Signatures Database (MSigDB).

To complement this study, single-cell RNA data from human TNBC cells were analyzed. The processed single-cell RNAseq data from Wu et al. (31) were downloaded from CZ CELLxGENE at https://cellxgene.cziscience.com/collections/dea97145-f712-431c-a223-6b5f565f362a. The raw count from this study GSE173634 was accessible via the National Center for Biotechnology Information (NCBI) Gene Expression Omnibus (GEO) database (https://www.ncbi.nlm.nih.gov/geo/). Only cancer cells originating from the TNBC patient whose cancer cells were PDPN-positive were used in our analysis, with patient ID 4513. Data normalization and dimensionality reduction, using default parameters, were conducted using scanpy (v.1.9.6). More specifically, dimensionality reduction was performed using Uniform Manifold Approximation and Projection (UMAP), and the clustering was conducted using Leiden algorithm with a resolution of 0.1. The Wilcoxon rank test was applied to identify DEGs between clusters, and significant genes were identified with Bonferroni-Hochberg corrected P value < 0.05 and |log2FC| > 1. Pathway analysis was performed using GSEApy (v.1.1.1) with human hallmark gene sets (v2023.2) obtained from the MSigDB.

### Mice and tumor inoculation

Female C57BL/6J mice were housed under specific pathogen–free (SPF) conditions. Mice aged 8-20 weeks were used for the experiments. All animal procedures were approved by the Uppsala County Regional Ethics Committee (Dnr: 6009/17 MHU and 5.8.18-03169.2023 MHU). For tumor inoculation, 4 × 10^5 cells in 5 μL PBS (Gibco) were injected into the 4th mammary fat pad of female mice. Tumor-bearing mice were sacrificed at a defined experimental endpoint or when tumors reached a maximum diameter of 12 mm or based on experimental setup, determined by caliper measurements. To assess tissue hypoxia, the Hypoxyprobe-PAb27 Kit (Hypoxyprobe Inc.) was used following the manufacturer’s instructions. Mice were intraperitoneally (i.p.) injected with pimonidazole hydrochloride at a dose of 60 mg/kg body weight, prepared in sterile PBS. The injection was administered 60 minutes before euthanasia. Tissues were fixed in 0.8% PFA overnight and processed by cryo-sectioning. Hypoxia was detected using the PAb27 monoclonal antibody from the kit, followed by immunofluorescence staining.

### Flow cytometry

LNs and tumor digestion were performed as previously described (45). The dead cells, RBCs (Ter119+) were gated away. Data were acquired on a BD FACSAriaIII flow cytometer (BD Biosciences) or Cytoflex and analyzed using FlowJo software (FlowJo, LLC, version 10.6.1). Antibodies are listed in Supplementary Table 3.

### Tissue Preparation and imaging

Organs for imaging were directly fixed in 0.8% paraformaldehyde (PFA) overnight at 4°C. For frozen section preparation, tissues were subsequently transferred to PBS, immersed in 25% sucrose, then 50% sucrose, and finally embedded in Optimal Cutting Temperature (OCT) compound. Samples were snap-frozen on dry ice and stored at −80°C until further use. LN sections were cut at 8–12 µm thickness using a cryostat, allowed to air dry 1h or overnight, and then stored at −80°C. Alternatively, after fixation, tissues were embedded into agarose and vibratome sections, 80μm were prepared. The PFA-fixed tissue was permeabilized in PBS. Samples were permeabilized in PBS with 0,1% Triton-X-100 (PBST). When required, endogenous biotin was blocked by treatment with 0.003% avidin in PBS containing (PBST), followed by 0.0033% biotin in PBST. Non-specific binding sites were blocked using 10% donkey serum in PBST. Sections were incubated with primary antibodies in blocking buffer overnight at 4°C or for 1 h at room temperature (RT). After washing with PBST, the samples were incubated with fluorophore-conjugated secondary antibodies in PBST for 45 min at RT, followed by additional washes in PBST. Nuclear staining was performed using 4′,6-diamidino-2-phenylindole (DAPI, Invitrogen) at a dilution of 1:5000. Slides were mounted using Prolong™ Gold Antifade Mounting Media (Thermo Fisher Scientific). The antibodies used are listed in Supplementary Table 3. Images were acquired using an automated slide scanner, Vectra Polaris (Akoya). Images for publishing were prepared using the open software Fiji^TM^ v.2.11.0 Software (ImageJ 2).

### Image analysis

Intratumoral T-cells: Image analysis was performed using Fiji sofware (ImageJ 2.11.0). Sections from the middle of each tumor were selected. Equally sized Regions of Interest (ROIs; 0.21 mm²) were randomly chosen, excluding tumor margins and necrotic areas. CD3+ CD8+ cells were manually counted in each frame. CD11b/MHC classII: The same ROI selection strategy as for T-cells was also applied to analyze CD11b+ myeloid cells and MHC class II+ cells, but cell density was determined instead of manual counting. A macro was used to ensure consistent analysis across all the images.

### METABRIC gene expression analyses

To explore CCR7-PDPN mRNA expression score, we analyzed publicly available gene expression datasets from primary BC with clinico-pathological data and follow-up information: Molecular Taxonomy of Breast Cancer International Consortium (METABRIC), METABRIC discovery (n=939) and validation cohort (nC=C845) (32). Information on intrinsic molecular subtypes based on the PAM50 classification was available (46). The normal-like category was excluded. JExpress/2012 (www.molmine.com) was used to extract differentially expressed genes from the METABRIC cohort. In cases of multiple probes per gene symbol in the gene expressions matrices, the probes were collapsed according to maximum probe expression per gene (47).

### Statistical methods METABRIC

Publicly available breast cancer gene expression data were analyzed using SPSS (version 25.0, IBM Corp., Armonk, NY, USA). Spearman’s rank correlation test was applied to compare bivariate continuous variables, and Spearman’s correlation coefficients (ρ) were reported. Mann–Whitney U or Kruskal–Wallis tests were applied to analyze the differences in the distribution of continuous variables between two or more categories. For univariate survival analyses, including death from breast cancer or recurrence from BC as endpoints, the Kaplan–Meier product-limit method (log-rank test for differences) was applied. In multivariate logistic regression analysis, the calculations were done according to the Backward Elimination (Likelihood Ration), with p-values derived from Step1 in the “model if term removed”-table. Moreover, only patients with information on all the variables were included. All statistical tests were two-sided and statistical significance was assessed at 5% level. CCR7-PDPN score dichotomization; quartile 1-3 = low CCR7-PDPN score; quartile 4 = high CCR7-PDPN score.

### Data availability statement

RNA-seq raw data from EO771 will be deposited to the NCBI Gene Expression Omnibus database upon publication, access numbers GSE-x. Code is available from https://github.com/zhi-igp-uu-se/ccr7andpdpn.git. METABRIC gene expression datasets are available at https://ega-archive.org/studies/EGAS00000000083 (Molecular Taxonomy of Breast Cancer International Consortium - METABRIC). Restrictions apply to data generated within this study—these are therefore not publicly available. However, on reasonable request, interested researchers may contact E.W and E.A.H to inquire about access. Request for noncommercial use will be considered and will require full ethics review

## Author contributions statement

MHU conceived of the project. MHU wrote the original manuscript with contribution of LMI and ZW. MHU with contribution of LMI, ZW, WS and TB designed the figures. ZW performed all bioinformatical analysis and contributed to experimental work, LMI performed METABRIC analysis, EW and EAH gave advice and contributed with data for the METABRIC analysis. AB, WS, TB, JHL, MV, AP, SR performed experimental work and analysis. SR and TB contributed to supervision and TB design of image analysis. JF contributed with advice and RNAseq. JF, EW, EAH and TB contributed to editing and improvement of the manuscript. All authors agreed to the manuscript.

## Acknowledgments

The authors thank Eira Lindqvist, Anton Johansson, Alexandra Rafeletou, Jennifer Mehjabin, and Kai Tang Xin for technical help and pilot studies. BioVis facility, Uppsala University, is acknowledged for flow cytometry and microscopy access. The authors also acknowledge the National bioinformatics infrastructure (NBIS) SciLifeLab Uppsala University for Bioinformatic support and training.

## Funding

This research was funded by the, Swedish Cancer Foundation (#200970 PjF and #232853 Pj), Kjell and Märta Beijer Foundation, and PO Zetterling Foundation to Ulvmar MH, Lise Martine Ingebriktsen was funded by Wennergren visiting researcher scholarship. Jonas Fuxe was funded by the Swedish Cancer Society (#21 1739 Pj, #24 3842 Pj), Radiumhemmet Research Funds (#211092, #231172 to J.F), and Karolinska Institutet Research Funds (#2024-03053 to J.F.).

## Conflict of Interest declaration

The authors declare that they have no affiliations with or involvement in any organization or entity with any financial interest in the subject matter or materials discussed in this manuscript.

## Supplementary Figures

**Suppl. Fig. 1: FACS CCR7 in EO771 with CCR7 expression and detection of tumor-derived tdTomato in LN macrophages.** A) Detection of surface expressed CCR7 by flow cytometry in control (EO771 X2 control vector, black and EO771 CCR7, dark gray). Y-axis normalized to mode. B) Staining of tdTomato (cyan) and CD169 (magenta) subcapsular macrophages, arrows indicate double staining. Scale bar 50 µm (representative of at least 6 TDLNs)

**Suppl. Fig. 2: *In vitro* characterization EO771 with and without CCR7 and PDPN and tumor take *in vivo*.** A) Histogram displaying the average tdTomato expression across the three different cell derivates. B) CCR7 expression is detected in both CCR7 PDPN positive and negative (green and orange) but not X2 PDPN negative cells. C) In vitro proliferation assay, based on CyQUANT® Cell Proliferation Assay Kit (Invitrogen). Data points indicate the mean values of three replicates per sample. The errors bars show Standard deviation (SD). Two-way Anova show no significant differences between groups). Y-axis normalized to mode. D) Tumor uptake across 30 injections in each group (15 mice) 50%, 70% and 80% X2, CCR7, and CCR7-PDPN respectively.

**Suppl. Fig. 3: GC formation and tdTomato-positive immunocomplex deposition on FDCs do not differ between EO771 with and without CCR7 and PDPN.** A) TDLNs from EO771 CCR7 stained for FDCs (CD21/35), follicular B-cells (IgD), tdTomato and nuclei (DAPI). Scale bar 50 µm. Stars indicate initiated GC formation based on downregulation of IgD in the center of the B-cell follicle. B-E) Analysis of TDLNs of end-stage tumors. Analysis of the TDLNs was performed in a central section, and consistency across samples is shown by equal number of follicles. B) TDLN size in mg. C) Numbers of follicles in a central section. D) Number of germinal centers (GC). E) tdTomato positive GCs. F-J) Analysis of TDLNs day 14 after tumor injection. F) Size of tumors in the groups used for analysis. G) Weight of TDLNs day 14 after tumor injection. H) Number of GCs. I) Analysis FDC (CD21/35) positive area in GC. X2 (n=11), CCR7 (n=12) and CCR7-PDPN (n=13). J). Integrated density for the tdTomato signal was calculated by tdTomato mean gray value x tdTomato area for each GC. No significant differences across groups. B-J) Kruskal Wallis Test and Dunn multiple comparison were used.

**Suppl. Fig. 4: EO771 X2 PDPN characterization.** A) Growth curve of X2 and X2 PDPN injected on both sides 4^th^ mammary fat pad, n=7 each group. Mice were followed over 14 days. No significant differences. Statistical evaluation using Two-way ANOVA. B-C) X2, CCR7 and CCR7-PDPN same groups as shown in Fig. 2, complemented with X2 PDPN tumors. B) Time after tumor inoculation for samples used for determination of metastasis in C). C) Metastasis was determined by manual assessment of at least 3 levels of each LN (C-D) X2 (n = 9), X2 PDPN (n=8), CCR7 (n = 8), CCR7-PDPN (n = 9).

**Suppl. Fig. 5: Colony forming assay EO771.** Representative images of colony on the left. 100 single cells were seeded in each well of 6-well plate and were allowed to grow for 14 days. Quantification of colony numbers is presented in the scatter plot with standard error of the mean (SEM) on the right. No significant differences were observed among the groups (n=3) using Kruskal-Wallis test with Dunn’s multiple comparison test.

**Suppl. Fig. 6: Hypoxyprobe staining.** Staining of an EO771-CCR7 tumor for hypoxi-probe (green) together with detection of tTomato (cyan), PDPN (magenta) Nuclei (DAPI grey), square indicate area shown in higher zoom in Fig. 3H. Single channel images shown in grey. Scale bar 2 mm. PDPN is seen in the dense capsule-like rim around the tumor and induced in central part of the tumor. Hypoxyprobe stain areas between live and necrotic cells.

**Suppl. Fig. 7: Immune and vascular cell density EO771.** A) Total density of CD11b in X2 vector control and CCR7-PDPN. n=3 and 10 areas per tumor, total 30. B) Total density of MCH class II positive cells. n=3 each group and 10 areas per tumor, total 30. A-B) analysis include random areas excluding tumor border and necrotic areas. Samples were taken from same samples used for T-cell evaluation, with significant differences in CD8 T-cells. C) Representative images of CD11b (cyan) and MHC class II (magenta) in tumors from X2 vector control and CCR7-PDPN. Scale bar 200 μm. D) Flow cytometric analysis of total PECAM1 positive cells in X2 and CCR7-PDPN EO771. No significant differences, Mann Whitney test.

**Suppl. Fig. 8: Expression of Key Genes Associated with Human Breast Cancer Subtypes and EMT in EO771 Cells with and Without CCR7 and PDPN.** Heatmap show the expression of selected key genes associated with human breast cancer subtypes and epithelial-to-mesenchymal transition (EMT). Data are presented as normalized counts. Rows represent genes, and columns represent cell derivates (CCR7-positive, PDPN-positive, and corresponding negative controls).

**Suppl. Fig. 9: Single-cell dataset analysis of human breast cancer cell lines.** A) UMAP plot illustrating tumor clusters, colored by subtypes, including TNBC (triple-negative breast cancer), LA (luminal A), LB (luminal B), and basal-like (non-malignant). B) Density plot showing the expression of PDPN across the UMAP space, highlighting cell lines with heterogenous expression. C) Leiden clustering of TNBC cell lines, upper (CAL51) lower (MDAMB436) and heatmap across the UMAP space for PDPN expression, with corresponding cell counts added for information.

## Notes

### Competing Interest Statement

The authors have declared no competing interest.

## References

1. Harbeck N, Penault-Llorca F, Cortes J, Gnant M, Houssami N, Poortmans P, et al. Breast cancer. Nature Reviews Disease Primers. 2019;5(1):66.

2. Carter CL, Allen C, Henson DE. Relation of tumor size, lymph node status, and survival in 24,740 breast cancer cases. Cancer. 1989;63(1):181–7.

3. Galassi C, Chan TA, Vitale I, Galluzzi L. The hallmarks of cancer immune evasion. Cancer Cell. 2024;42(11):1825–63.

4. Bergers G, Fendt SM. The metabolism of cancer cells during metastasis. Nat Rev Cancer. 2021;21(3):162–80.

5. Karlsson MC, Gonzalez SF, Welin J, Fuxe J. Epithelial-mesenchymal transition in cancer metastasis through the lymphatic system. Molecular oncology. 2017;11(7):781–91.

6. de Visser KE, Joyce JA. The evolving tumor microenvironment: From cancer initiation to metastatic outgrowth. Cancer Cell. 2023;41(3):374–403.

7. du Bois H, Heim TA, Lund AW. Tumor-draining lymph nodes: At the crossroads of metastasis and immunity. Sci Immunol. 2021;6(63):eabg3551.

8. Muller A, Homey B, Soto H, Ge N, Catron D, Buchanan ME, et al. Involvement of chemokine receptors in breast cancer metastasis. Nature. 2001;410(6824):50–6.

9. Cabioglu N, Yazici MS, Arun B, Broglio KR, Hortobagyi GN, Price JE, et al. CCR7 and CXCR4 as novel biomarkers predicting axillary lymph node metastasis in T1 breast cancer. Clin Cancer Res. 2005;11(16):5686–93.

10. Shields JD, Fleury ME, Yong C, Tomei AA, Randolph GJ, Swartz MA. Autologous chemotaxis as a mechanism of tumor cell homing to lymphatics via interstitial flow and autocrine CCR7 signaling. Cancer Cell. 2007;11(6):526–38.

11. Liu Y, Ji R, Li J, Gu Q, Zhao X, Sun T, et al. Correlation effect of EGFR and CXCR4 and CCR7 chemokine receptors in predicting breast cancer metastasis and prognosis. Journal of Experimental & Clinical Cancer Research. 2010;29(1):16.

12. Cunningham HD, Shannon LA, Calloway PA, Fassold BC, Dunwiddie I, Vielhauer G, et al. Expression of the C-C chemokine receptor 7 mediates metastasis of breast cancer to the lymph nodes in mice. Transl Oncol. 2010;3(6):354–61.

13. Ohl L, Mohaupt M, Czeloth N, Hintzen G, Kiafard Z, Zwirner J, et al. CCR7 governs skin dendritic cell migration under inflammatory and steady-state conditions. Immunity. 2004;21(2):279–88.

14. Forster R, Schubel A, Breitfeld D, Kremmer E, Renner-Muller I, Wolf E, et al. CCR7 coordinates the primary immune response by establishing functional microenvironments in secondary lymphoid organs. Cell. 1999;99(1):23–33.

15. Pang MF, Georgoudaki AM, Lambut L, Johansson J, Tabor V, Hagikura K, et al. TGF-beta1-induced EMT promotes targeted migration of breast cancer cells through the lymphatic system by the activation of CCR7/CCL21-mediated chemotaxis. Oncogene. 2016;35(6):748–60.

16. Sonbul SN, Gorringe KL, Aleskandarany MA, Mukherjee A, Green AR, Ellis IO, et al. Chemokine (C-C motif) receptor 7 (CCR7) associates with the tumour immune microenvironment but not progression in invasive breast carcinoma. The Journal of Pathology: Clinical Research. 2017;3(2):105–14.

17. Weitzenfeld P, Kossover O, Korner C, Meshel T, Wiemann S, Seliktar D, et al. Chemokine axes in breast cancer: factors of the tumor microenvironment reshape the CCR7-driven metastatic spread of luminal-A breast tumors. J Leukoc Biol. 2016;99(6):1009–25.

18. Johnstone CN, Smith YE, Cao Y, Burrows AD, Cross RS, Ling X, et al. Functional and molecular characterisation of EO771.LMB tumours, a new C57BL/6-mouse-derived model of spontaneously metastatic mammary cancer. Dis Model Mech. 2015;8(3):237–51.

19. Le Naour A, Rossary A, Vasson M-P. EO771, is it a well-characterized cell line for mouse mammary cancer model? Limit and uncertainty. Cancer medicine. 2020;9(21):8074–85.

20. Crosby EJ, Wei J, Yang XY, Lei G, Wang T, Liu CX, et al. Complimentary mechanisms of dual checkpoint blockade expand unique T-cell repertoires and activate adaptive anti-tumor immunity in triple-negative breast tumors. Oncoimmunology. 2018;7(5):e1421891.

21. Gu Y, Zhang Z, ten Dijke P. Harnessing epithelial-mesenchymal plasticity to boost cancer immunotherapy. Cellular & Molecular Immunology. 2023;20(4):318–40.

22. Ye X, Weinberg RA. Epithelial-Mesenchymal Plasticity: A Central Regulator of Cancer Progression. Trends in cell biology. 2015;25(11):675–86.

23. Bill CA, Allen CM, Vines CM. C-C Chemokine Receptor 7 in Cancer. Cells. 2022;11(4):656.

24. Quintanilla M, Montero-Montero L, Renart J, Martin-Villar E. Podoplanin in Inflammation and Cancer. International journal of molecular sciences. 2019;20(3).

25. Breiteneder-Geleff S, Soleiman A, Kowalski H, Horvat R, Amann G, Kriehuber E, et al. Angiosarcomas express mixed endothelial phenotypes of blood and lymphatic capillaries: podoplanin as a specific marker for lymphatic endothelium. Am J Pathol. 1999;154(2):385–94.

26. de Winde CM, George SL, Crosas-Molist E, Hari-Gupta Y, Arp AB, Benjamin AC, et al. Podoplanin drives dedifferentiation and amoeboid invasion of melanoma. iScience. 2021;24(9):102976.

27. Bresson L, Faraldo MM, Di-Cicco A, Quintanilla M, Glukhova MA, Deugnier MA. Podoplanin regulates mammary stem cell function and tumorigenesis by potentiating Wnt/beta-catenin signaling. Development. 2018;145(4).

28. Heesters BA, Myers RC, Carroll MC. Follicular dendritic cells: dynamic antigen libraries. Nat Rev Immunol. 2014;14(7):495–504.

29. Chen DS, Mellman I. Oncology meets immunology: the cancer-immunity cycle. Immunity. 2013;39(1):1–10.

30. Gambardella G, Viscido G, Tumaini B, Isacchi A, Bosotti R, di Bernardo D. A single-cell analysis of breast cancer cell lines to study tumour heterogeneity and drug response. Nat Commun. 2022;13(1):1714.

31. Wu SZ, Al-Eryani G, Roden DL, Junankar S, Harvey K, Andersson A, et al. A single-cell and spatially resolved atlas of human breast cancers. Nature Genetics. 2021;53(9):1334–47.

32. Curtis C, Shah SP, Chin SF, Turashvili G, Rueda OM, Dunning MJ, et al. The genomic and transcriptomic architecture of 2,000 breast tumours reveals novel subgroups. Nature. 2012;486(7403):346-52.

33. Sasano T, Gonzalez-Delgado R, Munoz NM, Carlos-Alcade W, Cho MS, Sheth RA, et al. Podoplanin promotes tumor growth, platelet aggregation, and venous thrombosis in murine models of ovarian cancer. J Thromb Haemost. 2022;20(1):104–14.

34. Li X, Jin Y, Xue J. Unveiling Collagen’s Role in Breast Cancer: Insights into Expression Patterns, Functions and Clinical Implications. Int J Gen Med. 2024;17:1773–87.

35. Papalazarou V, Drew J, Juin A, Spence HJ, Whitelaw J, Nixon C, et al. Collagen VI expression is negatively mechanosensitive in pancreatic cancer cells and supports the metastatic niche. J Cell Sci. 2022;135(24).

36. Lei PJ, Pereira ER, Andersson P, Amoozgar Z, Van Wijnbergen JW, O’Melia MJ, et al. Cancer cell plasticity and MHC-II-mediated immune tolerance promote breast cancer metastasis to lymph nodes. J Exp Med. 2023;220(9).

37. Gozgit JM, Vasbinder MM, Abo RP, Kunii K, Kuplast-Barr KG, Gui B, et al. PARP7 negatively regulates the type I interferon response in cancer cells and its inhibition triggers antitumor immunity. Cancer Cell. 2021;39(9):1214–26.e10.

38. Kwon J, Bakhoum SF. The Cytosolic DNA-Sensing cGAS–STING Pathway in Cancer. Cancer Discovery. 2020;10(1):26–39.

39. Hu J, Sánchez-Rivera FJ, Wang Z, Johnson GN, Ho YJ, Ganesh K, et al. STING inhibits the reactivation of dormant metastasis in lung adenocarcinoma. Nature. 2023;616(7958):806–13.

40. Rasmussen M, Alvik K, Kannen V, Olafsen NE, Erlingsson LAM, Grimaldi G, et al. Loss of PARP7 Increases Type I Interferon Signaling in EO771 Breast Cancer Cells and Prevents Mammary Tumor Growth by Increasing Antitumor Immunity. Cancers (Basel). 2023;15(14).

41. Louie DAP, Oo D, Leung G, Lin Y, Stephens M, Alrashed O, et al. Tumor-Draining Lymph Node Reconstruction Promotes B Cell Activation During E0771 Mouse Breast Cancer Growth. Frontiers in pharmacology. 2022;13:825287.

42. Reddy TP, Rosato RR, Li X, Moulder S, Piwnica-Worms H, Chang JC. A comprehensive overview of metaplastic breast cancer: clinical features and molecular aberrations. Breast Cancer Research. 2020;22(1):121.

43. Lien H-C, Hsu C-L, Lu Y-S, Chen TW-W, Chen IC, Li Y-C, et al. Transcriptomic alterations underlying metaplasia into specific metaplastic components in metaplastic breast carcinoma. Breast Cancer Research. 2023;25(1):11.

44. Sarrio D, Franklin CK, Mackay A, Reis-Filho JS, Isacke CM. Epithelial and mesenchymal subpopulations within normal basal breast cell lines exhibit distinct stem cell/progenitor properties. Stem Cells. 2012;30(2):292–303.

45. Xiang M, Grosso RA, Takeda A, Pan J, Bekkhus T, Brulois K, et al. A Single-Cell Transcriptional Roadmap of the Mouse and Human Lymph Node Lymphatic Vasculature. Frontiers in Cardiovascular Medicine. 2020;7(52).

46. Sørlie T, Perou CM, Tibshirani R, Aas T, Geisler S, Johnsen H, et al. Gene expression patterns of breast carcinomas distinguish tumor subclasses with clinical implications. Proc Natl Acad Sci U S A. 2001;98(19):10869–74.

47. Subramanian A, Tamayo P, Mootha VK, Mukherjee S, Ebert BL, Gillette MA, et al. Gene set enrichment analysis: a knowledge-based approach for interpreting genome-wide expression profiles. Proc Natl Acad Sci U S A. 2005;102(43):15545–50.

