## Supplementary figures and images for "Podoplanin-Linked Mesenchymal Shift Synergizes with CCR7 driving Lymphatic Metastasis and Tumor Progression in Breast Cancer"

### S1

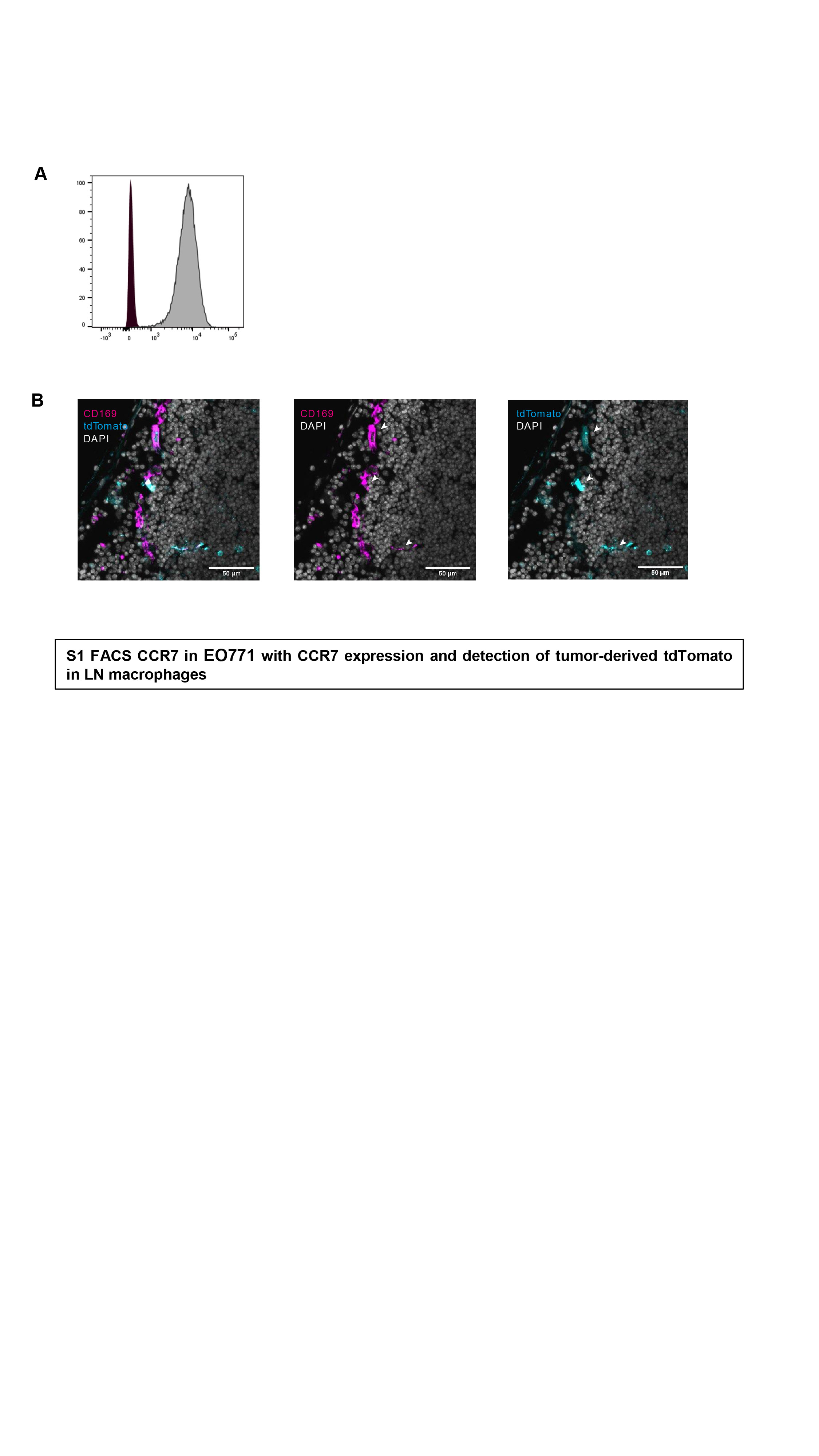

### S2

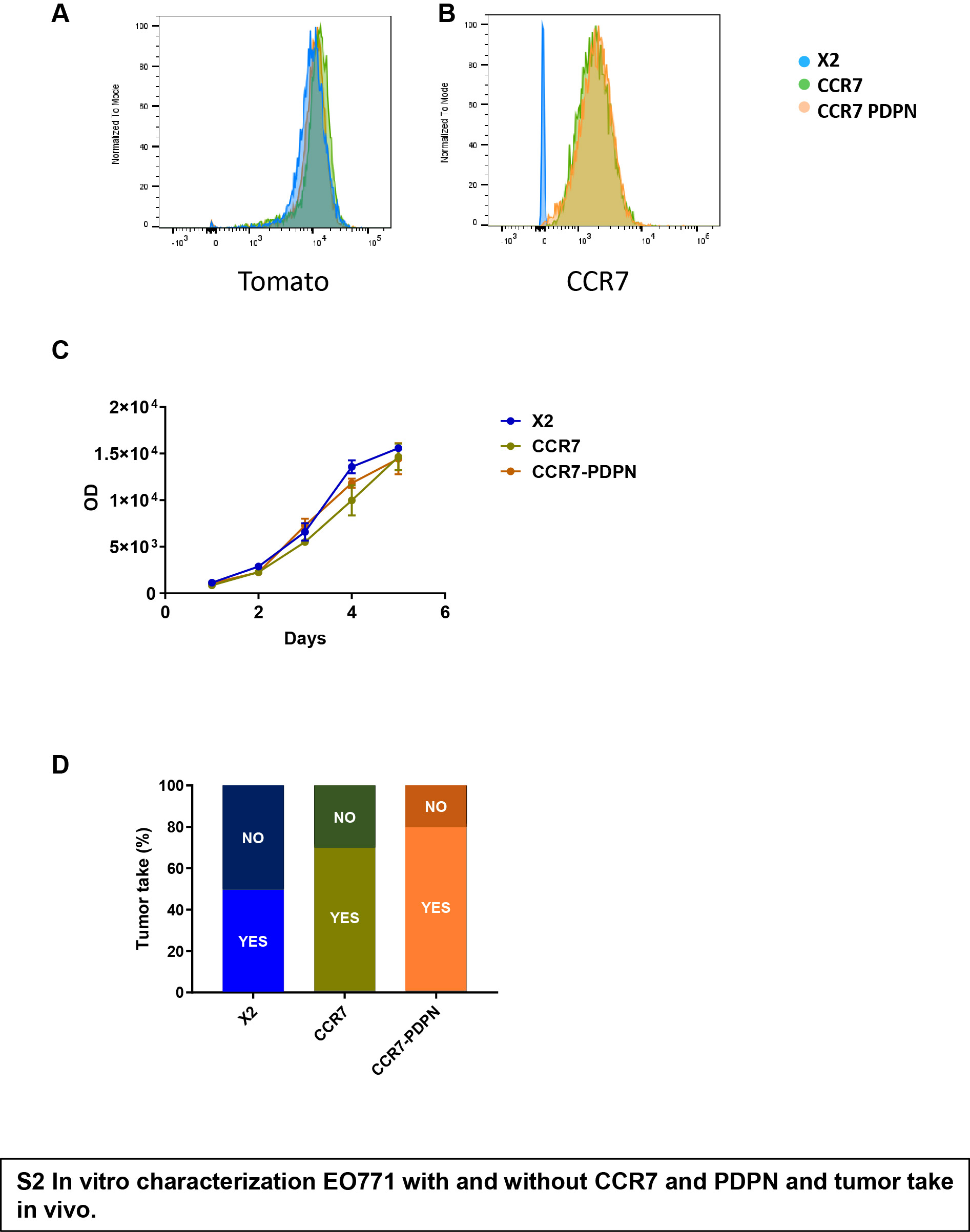

### S3

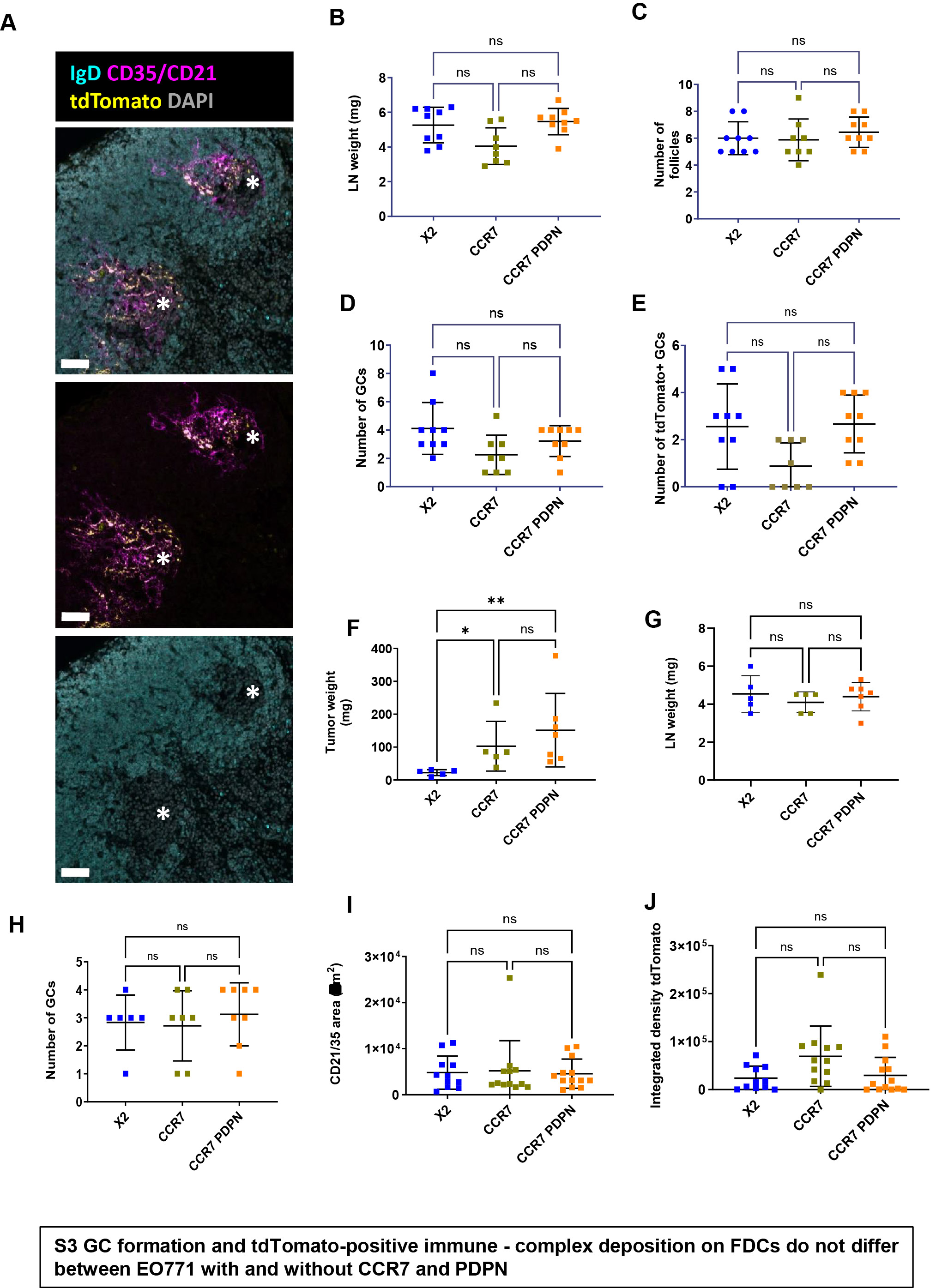

### S4

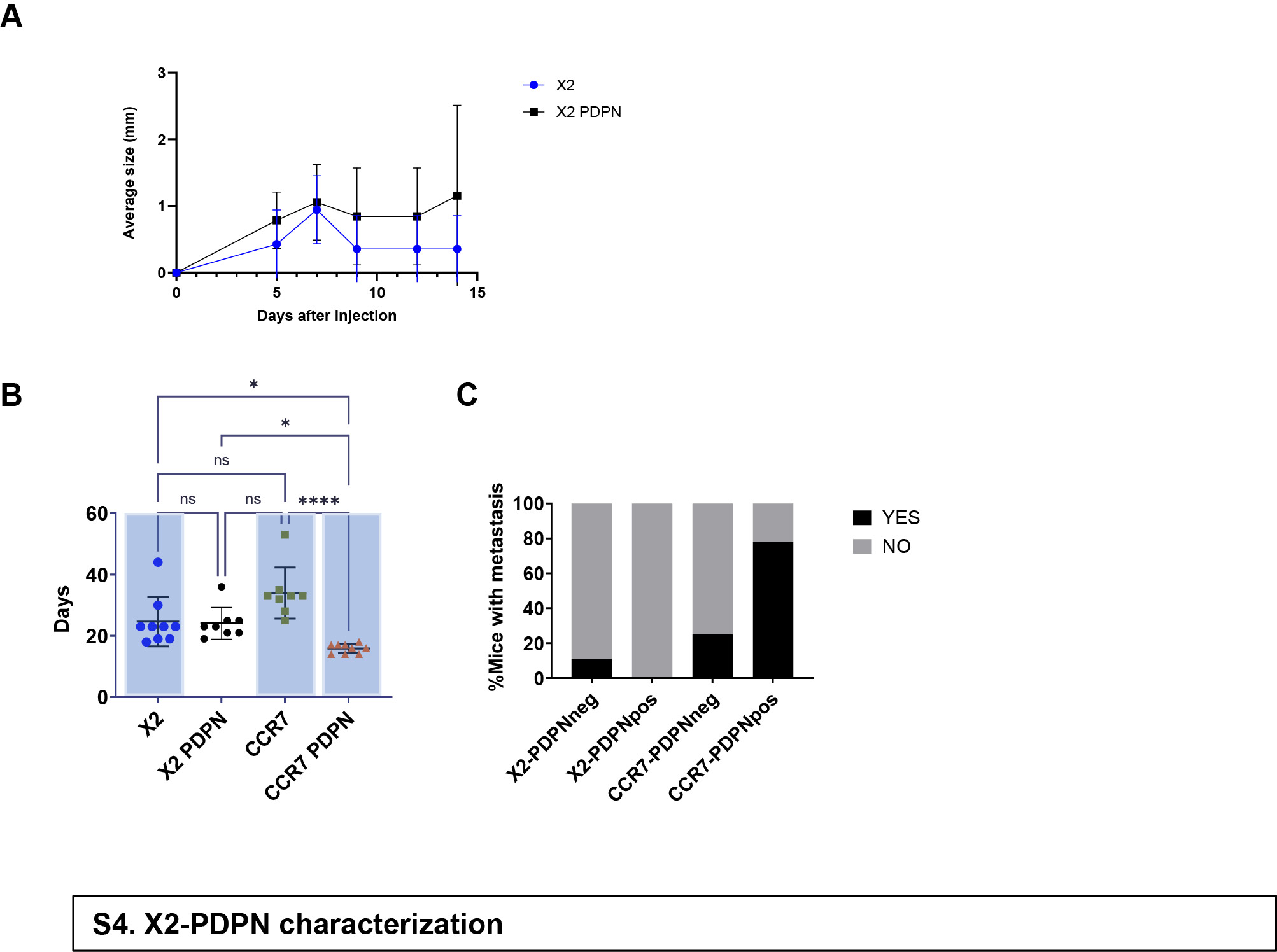

### S5

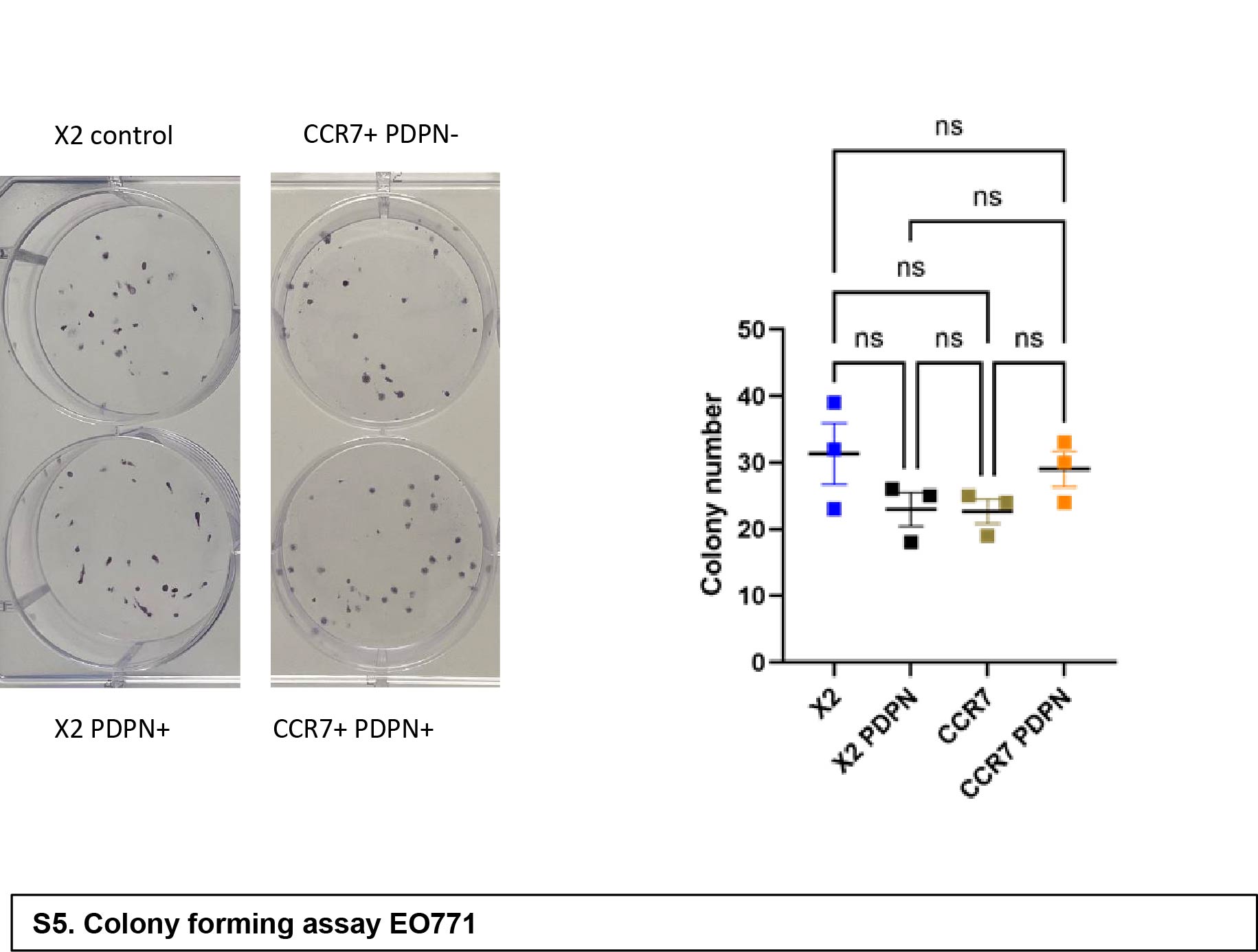

### S6

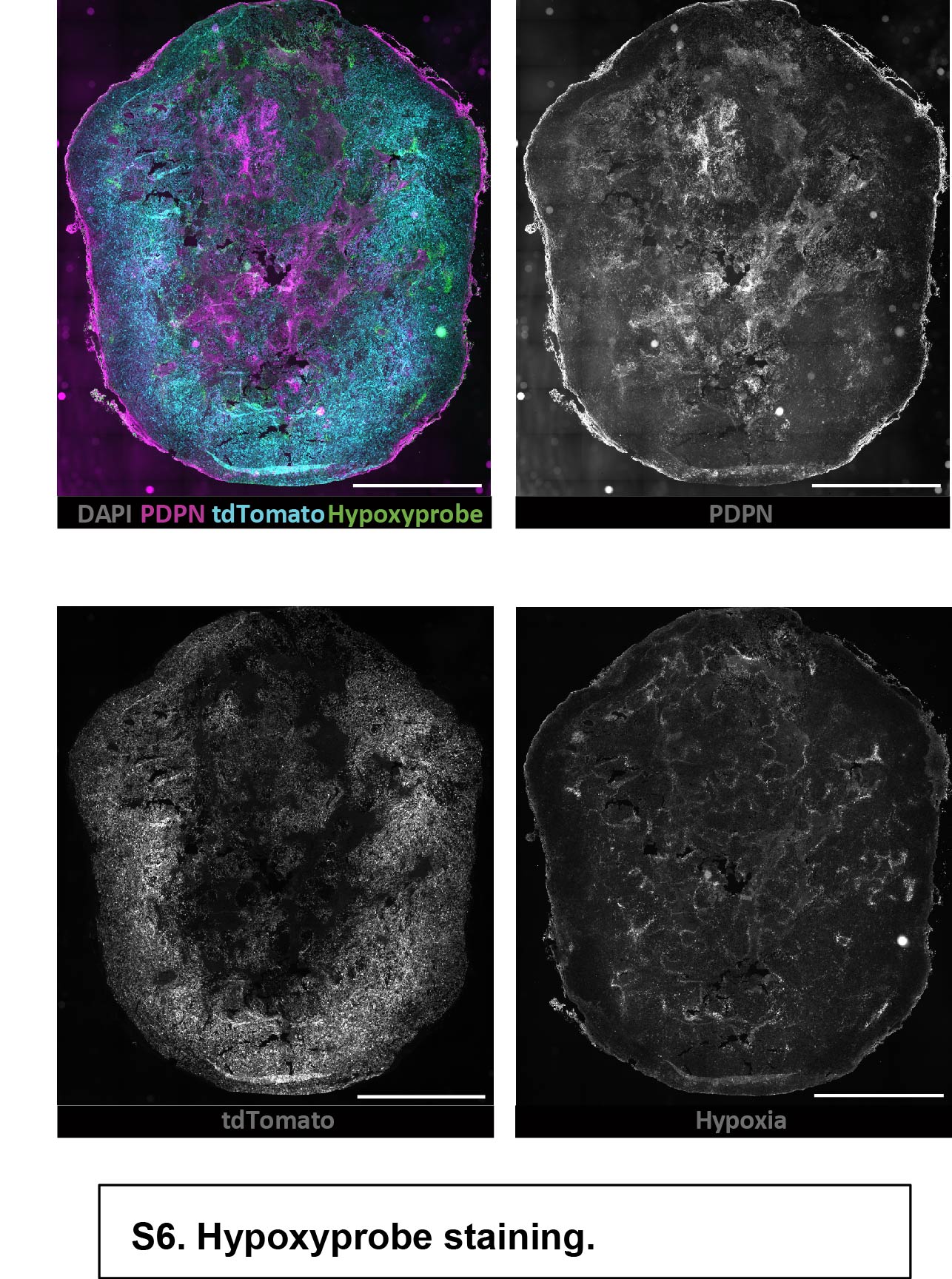

### S7

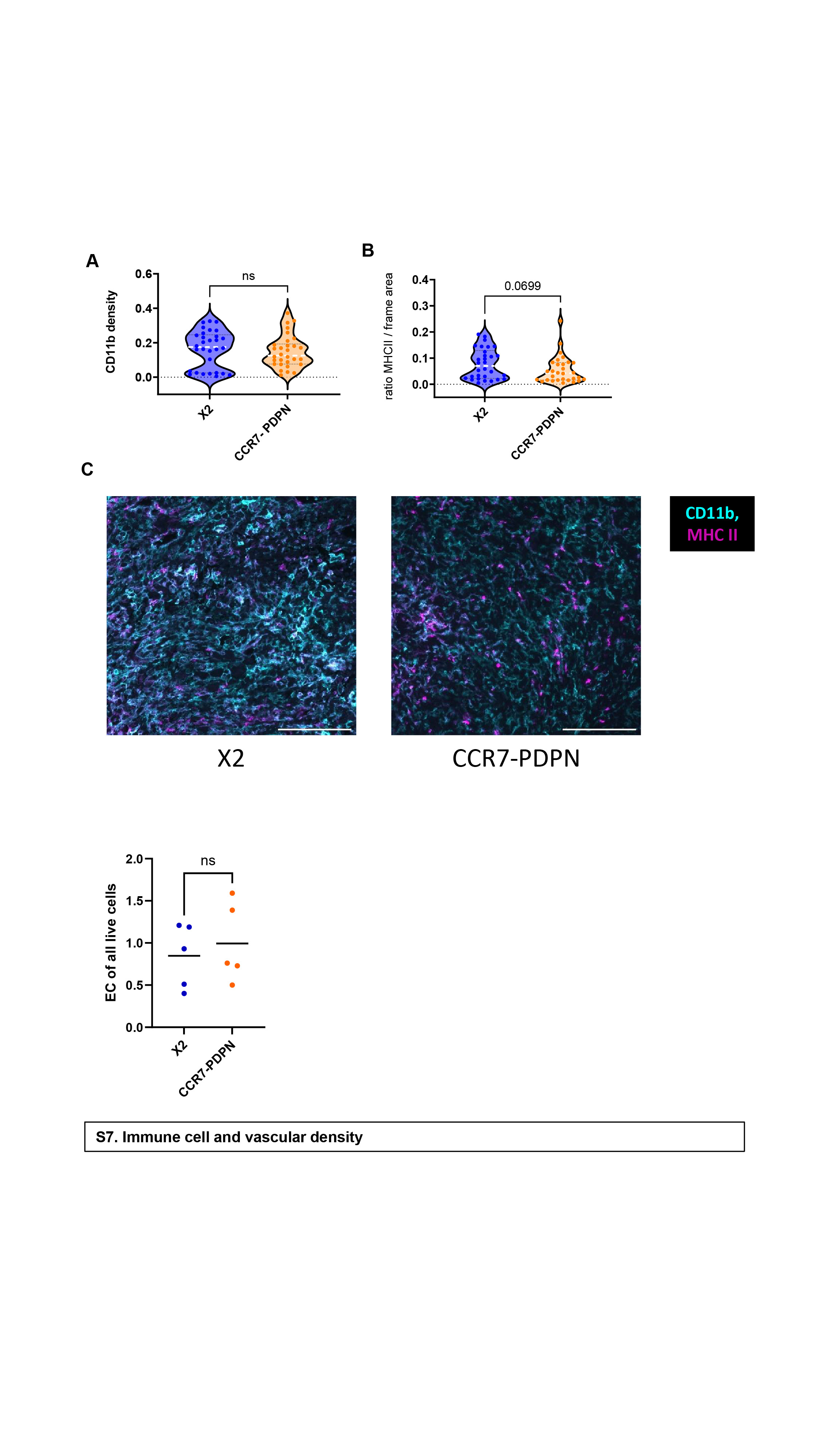

### S8

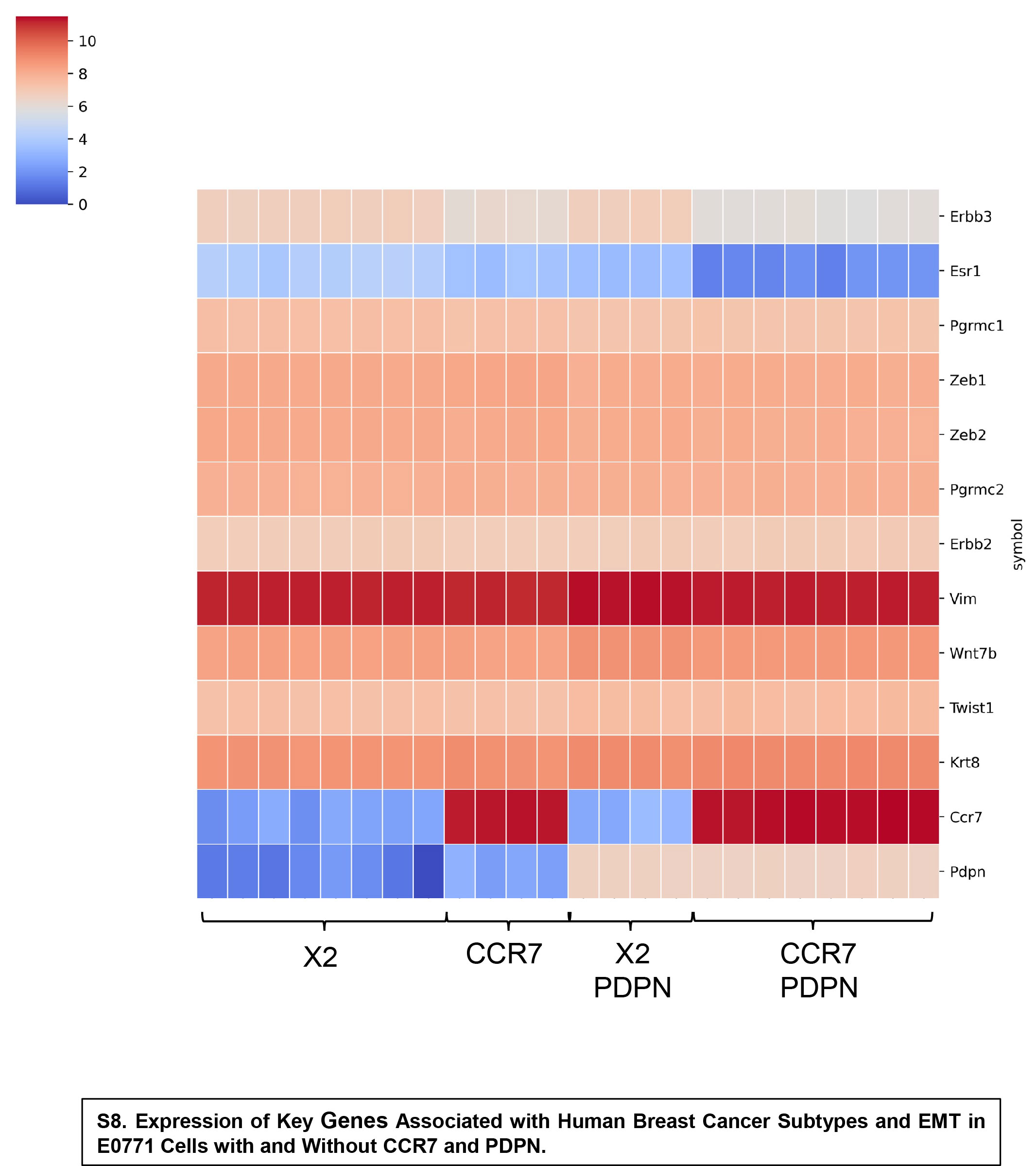

### S9

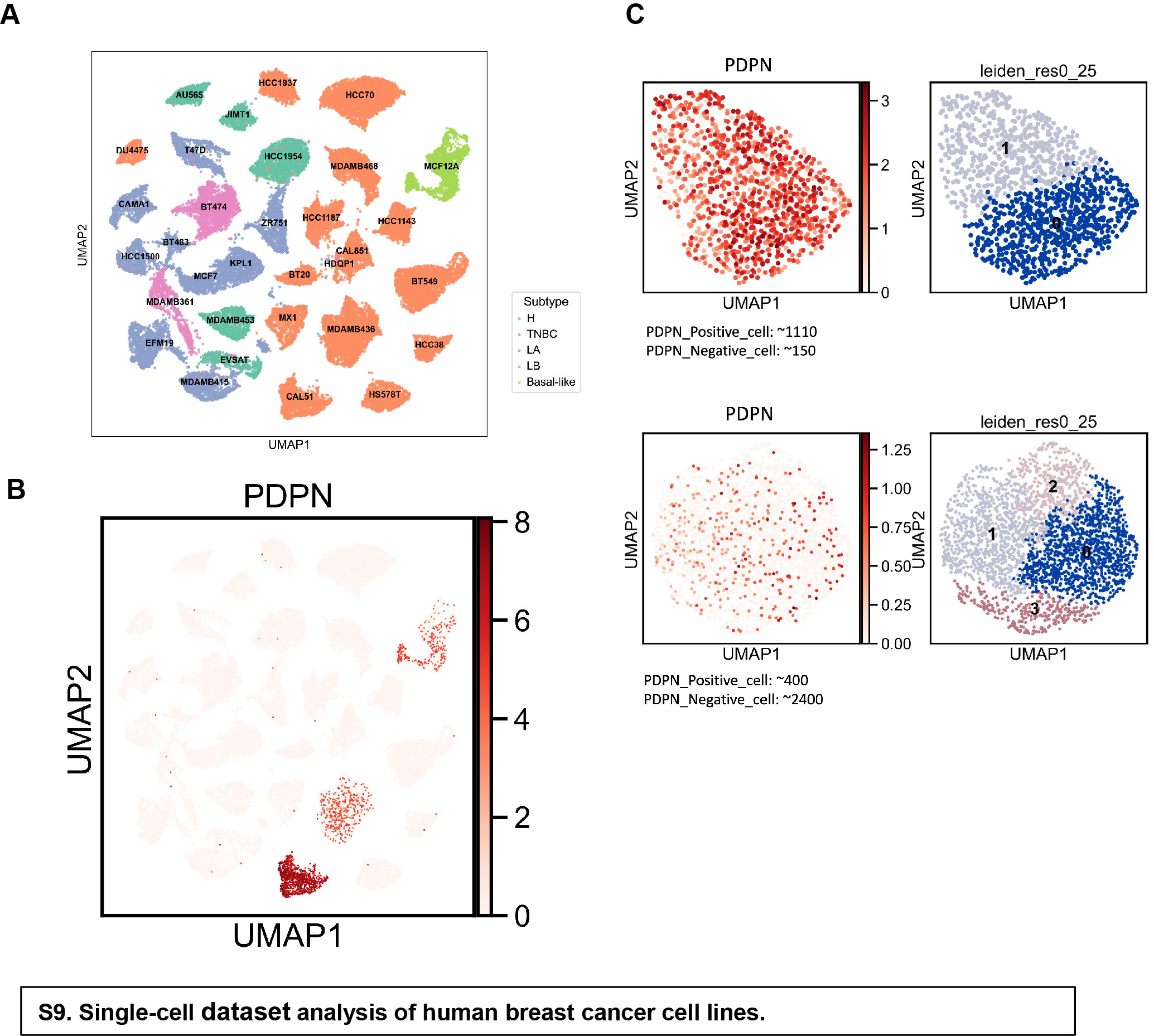
