## Supplementary material for "Podoplanin-Linked Mesenchymal Shift Synergizes with CCR7 driving Lymphatic Metastasis and Tumor Progression in Breast Cancer": S Table 2

**Supplementary Table 2 A-B: Univariate and multivariate survival analyses** (Cox' proportional hazards regression) with death from breast cancer as end-point. All subtypes (**A**: METABRIC validation cohort, n=845) and all subtypes, lymph node metastasis only (**B**: METABRIC validation cohort, n=405).
 **2A**

| **METABRIC validation cohort, All subtypes** | | | | | |
| --- | --- | --- | --- | --- | --- |
| **Variables** | **n** | **Univariate HR (95% CI)** | **p** | **Multivariate (95% CI)** | **p** |
| **Tumor diameter** |  |  |  |  |  |
| <2.0 cm | 347 | 1 |  | 1 |  |
| ≥2.0 cm | 498 | 2.00 (1.49-2.67) | **<0.001** | 1.94 (1.42-2.63) | **<0.001** |
| **Histologic grade*** |  |  |  |  |  |
| 1+2 | 363 | 1 |  | 1 |  |
| 3 | 409 | 1.66 (1.25-2.20) | **<0.001** | 1.23 (0.89-1.68) | **NS** |
| **Nodal status** |  |  |  |  |  |
| Negative | 440 | 1 |  | 1 |  |
| Positive | 405 | 2.86 (2.14-3.80) | **<0.001** | 2.34 (1.73-3.15) | **<0.001** |
| **ER status**** |  |  |  |  |  |
| ER positivity | 586 | 1 |  | 1 |  |
| ER negativity | 223 | 1.76 (1.33-2.32) | **<0.001** | 1.48 (1.10-2.00) | **0.008** |
| **CCR7-PDPN score***** |  |  |  |  |  |
| Low expression | 634 | 1 |  | 1 |  |
| High expression | 211 | 1.45 (1.09-1.93) | **0.010** | 1.40 (1.03-1.91) | **0.029** |

*Missing: 73 **Missing: 36 ***Cut-point Q1-3 low CCR7-PDPN score Q4 high CCR7-PDPN score
n= number of patients; HR: Hazard Ratio; CI: Confidence interval; P: p-values. NS: Not significant. Cox regression analysis (backward stepwise model)

**2B**

| **METABRIC validation cohort, All subtypes, LN met+ only** | | | | | |
| --- | --- | --- | --- | --- | --- |
| **Variables** | **n** | **Univariate HR (95% CI)** | **p** | **Multivariate (95% CI)** | **p** |
| **Tumor diameter** |  |  |  |  |  |
| <2.0 cm | 138 | 1 |  | 1 |  |
| ≥2.0 cm | 267 | 1.82 (1.26-2.62) | **0.001** | 1.97 (1.34-2.89) | **<0.001** |
| **Histologic grade*** |  |  |  |  |  |
| 1+2 | 162 | 1 |  | 1 |  |
| 3 | 220 | 1.48 (1.04-2.09) | **0.028** | 1.16 (0.79-1.70) | NS |
| **ER status**** |  |  |  |  |  |
| ER positivity | 279 | 1 |  | 1 |  |
| ER negativity | 124 | 1.45 (1.03-2.03) | **0.029** | 1.19 (0.81-1.74) | NS |
| **CCR7-PDPN score***** |  |  |  |  |  |
| Low expression | 302 | 1 |  | 1 |  |
| High expression | 103 | 1.70 (1.21-2.38) | **0.002** | 1.81 (1.28-2.58) | **<0.001** |

*Missing: 23 **Missing: 2 ***Cut-point Q1-3 low CCR7-PDPN score Q4 high CCR7-PDPN score
n= number of patients; HR: Hazard Ratio; CI: Confidence interval; P: p-values. NS: Not significant. LN Met+: Lymph node metastasis positive. Cox regression analysis (backward stepwise model)
