## Supplementary material for "Podoplanin-Linked Mesenchymal Shift Synergizes with CCR7 driving Lymphatic Metastasis and Tumor Progression in Breast Cancer": S Table 3

Supplementary Table 3: Overview Antibodies for Flow cytometry and FACS.

Flow cytometry

| **Antibody** | **Clone** | **Company** |
| --- | --- | --- |
| PerCP-Cy5.5 anti-CD11b | M1/70 | Thermofisher scientific |
| PerCP-Cy5.5 anti-CD45 | 30-F11 | Thermofisher scientific |
| BV421 anti-Ter119 | TER-119 | BD Biosciences |
| eFluor^TM^ 450 anti-Ter119 | TER-119 | Thermofisher scientific |
| eFluor^TM^ 660 anti-Podoplanin | eBioB.1.1 | Thermofisher scientific |
| PE-Cyanine7 anti-CD31 (PECAM-1) | 390 | Thermofisher scientific |
| APC anti-CCR7 | 4B12 | Thermofisher scientific |

Antibodies FACS

| **1*Antibodies** | **Clone** | **Company** | **Dilution** |
| --- | --- | --- | --- |
| Anti-mouse-IgD-FITC | 11-26c | Thermofisher scientific | 1:50 |
| Anti-mouse-IgM-FITC | 11/41 | Thermofisher scientific | 1:50 |
| tdTomato | Goat polyclonal | Sicgenantibodies | 1:200 |
| AF647- and FITC- anti-CD21/CD35 | 7E9 | BioLegend | 1:50 |
| Anti-LYVE1 | Rabbit polyclonal | ReliaTech | 1:200 |
| Anti-CD169 | HSn 7D2 | Santa Cruz Biotechnology | 1:100 |
| CD3e-FITC | eBio500A2 | Thermofisher scientific | 1:50 |
| Anti-PDGFRB | Rabbit 28E1 | Cell Signaling | 1:50 |
| anti-PDPN-eFluor™660 | eBioB.1.1 | ebiociences | 1:50 |
| Anti-CD8 | UCH-T4 | Santa Cruz Biotechnology | 1:50 |
| Anti-FITC-488 | Mouse Monoclonal | Jackson ImmunoResearch | 1:300 |

| **2* Antibodies** | **Clone** | **Company** | **Dilution** |
| --- | --- | --- | --- |
| anti-Rabbit AF647 | Polyclonal Donkey | Thermofisher scientific | 1:300 |
| anti-Goat AF555 | Polyclonal Donkey | Thermofisher scientific | 1:300 |
| anti-Rabbit AF488 | Polyclonal Donkey | Thermofisher scientific | 1:300 |
| anti-Rabbit AF488 | Polyclonal Donkey | Jackson ImmunoResearch | 1:300 |
| anti-Goat AF555 | Polyclonal Donkey | Thermofisher scientific | 1:300 |
| anti-Rat AF488 | Polyclonal Donkey | Jackson ImmunoResearch | 1:300 |
